# Subanesthetic ketamine reactivates adult cortical plasticity to restore vision from amblyopia

**DOI:** 10.1101/2020.03.16.994475

**Authors:** Steven F. Grieco, Xin Qiao, Xiaoting Zheng, Yongjun Liu, Lujia Chen, Hai Zhang, Jeffrey Gavornik, Cary Lai, Sunil Gandhi, Todd C. Holmes, Xiangmin Xu

**Author notes:** Lead contact / Address all manuscript correspondence to: Dr. Xiangmin Xu, Department of Anatomy and Neurobiology, School of Medicine, University of California, Irvine, CA 92697-1275. Data Availability Statement: The data that support the findings of this study are available from the corresponding author upon reasonable request. **Author contributions:** S.F.G performed molecular, viral, and behavioral experiments. X.Q. performed electrophysiology. Y.L. and H.Z. performed 2-photon calcium imaging; X.Z., and S.F.G performed intrinsic signal imaging; S.F.G and J.G. performed VEP experiments. S.P.G. supervised the intrinsic signal imaging and visual water maze experiments. L.C. analyzed calcium image data. X.X., S.F.G., T.C.H. and C.L. analyzed the data, prepared the figures and wrote the manuscript with the help and input from other authors. X.X. designed and oversaw the project.

## Abstract

Subanesthetic ketamine evokes rapid and long-lasting antidepressant effects in human patients. The mechanism for ketamine’s effects remains elusive, but ketamine may broadly modulate brain plasticity processes. We show that single-dose ketamine reactivates adult mouse visual cortical plasticity and promotes functional recovery of visual acuity defects from amblyopia. Ketamine specifically induces down-regulation of neuregulin-1 (NRG1) expression in parvalbumin-expressing (PV) inhibitory neurons in mouse visual cortex. NRG1 downregulation in PV neurons co-tracks both the fast onset and sustained decreases in synaptic inhibition to excitatory neurons, along with reduced synaptic excitation to PV neurons *in vitro* and *in vivo* following a single ketamine treatment. These effects are blocked by exogenous NRG1 as well as PV targeted receptor knockout. Thus ketamine reactivation of adult visual cortical plasticity is mediated through rapid and sustained cortical disinhibition via downregulation of PV-specific NRG1 signaling. Our findings reveal the neural plasticity-based mechanism for ketamine-mediated functional recovery from adult amblyopia.

**Highlights:**

○ Disinhibition of excitatory cells by ketamine occurs in a fast and sustained manner
○ Ketamine evokes NRG1 downregulation and excitatory input loss to PV cells
○ Ketamine induced plasticity is blocked by exogenous NRG1 or its receptor knockout
○ PV inhibitory cells are the initial functional locus underlying ketamine’s effects

## Introduction

Ketamine is a dissociative general anesthetic that acts as a non-competitive antagonist of N-methyl-D-aspartate receptors (NMDARs). Subanesthetic ketamine has been used as an effective antidepressant in patients who are resistant to typical antidepressants (Berman et al., 2000; Duman and Aghajanian, 2012). A single subanesthetic dose of ketamine evokes long lasting behavioral effects up to ∼2 weeks (Berman et al., 2000; Price et al., 2009; Zarate et al., 2006), extending well beyond ketamine’s chemical half-life time of ∼2 hours (Autry et al., 2011). While the molecular signaling and neural circuit mechanisms underlying ketamine’s acute and sustained effects remain to be elucidated, modulation of neural circuit plasticity is a potential explanation for these long lasting effects that exceed the persistence of ketamine or its biologically active metabolites.

Ketamine alters the neurochemical phenotype of parvalbumin-expressing (PV) inhibitory interneurons (Behrens et al., 2007). Ketamine and its metabolite, hydroxynorketamine (HNK) modulate cortical circuit excitation and inhibition (Li et al., 2010; Zanos et al., 2016). While it is controversial, it has been proposed that ketamine preferentially block NMDARs in inhibitory GABAergic neurons to reduce their excitatory input. This reduces inhibitory synaptic transmission onto excitatory glutamatergic neurons, thus promoting cortical disinhibition and increasing cortical excitability (Browne and Lucki, 2013; Homayoun and Moghaddam, 2007; Krystal et al., 2013). Although cortical disinhibition may initiate neural plasticity (Miller et al., 2016), the time course and the molecular mechanism remain unresolved. Previous studies identify signaling through several pathways implicated in ketamine’s action, including the release of brain-derived neurotrophic factor (BDNF) (Jourdi et al., 2009; Lepack et al., 2014), induction of structural plasticity by protein synthesis via eukaryotic elongation factor 2 (eEF2) deactivation, dendrite spine outgrowth by mammalian target of rapamycin (mTOR) (Moda-Sava et al., 2019; Takei et al., 2004; Xu et al., 2010; Zhou et al., 2014), and inhibition of glycogen synthase kinase-3 (GSK3) (Beurel et al., 2016; Beurel et al., 2011). But the mechanism for how ketamine works through neural activity-dependent molecular signaling in specific neuronal circuitry to induce cortical plasticity in a rapid and sustained fashion has not been determined.

Neuregulin-1 (NRG1) is essential for the normal development of the nervous system, and signaling through its tyrosine kinase receptor ErbB4 has been implicated in visual cortical plasticity (Sun et al., 2016; Gu et al., 2016; Grieco et al., 2019a). Our previous work demonstrates that PV inhibitory interneurons are the initial circuit locus for the establishment of visual cortex critical period plasticity (Kuhlman et al., 2013). While NRG1 is expressed in multiple cell types including excitatory neurons, inhibitory neurons and astrocytes (Liu et al., 2011), PV neurons have strong NRG1 expression relative to other cortical neurons, and the expression of NRG1’s cognate receptor ErbB4 is highly restricted to PV neurons in the visual cortex (Fazzari et al., 2010; Grieco et al., 2019b; Sun et al., 2016; Yang et al., 2013). In comparison, a relatively small proportion of vasoactive intestinal peptide (VIP) expressing inhibitory cells are immunopositve for ErbB4, and there is virtually no co-localization between somatostatin expressing inhibitory cells and ErbB4 (Sun et al., 2016). The co-expression of the ligand NRG1 and its receptor ErbB4 in PV neurons may allow these neurons to effectively regulate their synaptic plasticity through activity-dependent NRG1/ErbB4 signaling (Mei and Xiong, 2008). Monocular deprivation during the critical period down-regulates NRG1/ErbB4 signaling in PV neurons. This evokes rapid retraction of excitatory inputs to PV neurons, causing cortical disinhibition that is necessary for initiating critical period visual cortical plasticity (Sun et al., 2016). Based on these findings and other work showing that modulation of NRG1/ErbB4 signaling in PV neurons enhances adult visual cortex plasticity (Gu et al., 2016), we test the hypothesis that subanesthetic ketamine treatment reactivates adult visual cortical plasticity and promotes visual functional recovery by regulating NRG1-directed signaling in PV inhibitory neurons.

In the present study, we find that subanesthetic ketamine induces down-regulation of NRG1 expression in PV inhibitory neurons, resulting in PV excitatory input loss and sustained cortical disinhibition to enhance cortical plasticity in adult cortex. Our study establishes molecular, cellular, and circuit mechanisms of ketamine-mediated induction of adult cortical plasticity to promote functional recovery from amblyopia, which reopens the otherwise developmentally restricted critical period.

## Results

### Ketamine reactivates adult cortical plasticity

We first tested whether subanesthetic ketamine treatment effectively induces visual cortical plasticity during adulthood and well past the closure of the normal critical period for ocular dominance plasticity (Gordon and Stryker, 1996). Adult mice (P90-P120) were subjected to monocular deprivation for 4 days, with treatments of either saline or a single dose of subanesthetic ketamine (s.c.) immediately after the beginning of monocular deprivation via eyelid sutures (Figure 1A,B). Four days later, cortical plasticity measured by the ocular dominance index (ODI) of mice was assessed using intrinsic signal optical imaging of responses to visual stimuli (Davis et al., 2015; Kalatsky and Stryker, 2003; Southwell et al., 2010). Normally, four days of monocular deprivation does not induce visual cortical plasticity in adult control mice (Davis et al., 2015). In comparison, the mice receiving a single subanesthetic ketamine treatment exhibit significant visual cortical plasticity with 4 days of monocular deprivation (Figure 1C,D). Note that the selective reduction of deprived eye visual cortical responses associated with ketamine treatment, resembles the feature of visual cortical plasticity observed in critical period animals after monocular deprivation. Thus subanesthetic ketamine is effective in reactivating developmental critical period-like plasticity in the adult visual cortex. Further, we did not observe robust ketamine-induced visual cortical plasticity in PV-Cre; ErbB4^fl/fl^ mice, where ErbB4 is specifically ablated from PV interneurons (Long et al., 2003) (Figure S1). This supports the involvement of NRG1/ErbB4 signaling in ketamine-induced adult cortical plasticity.

**Figure 1.**
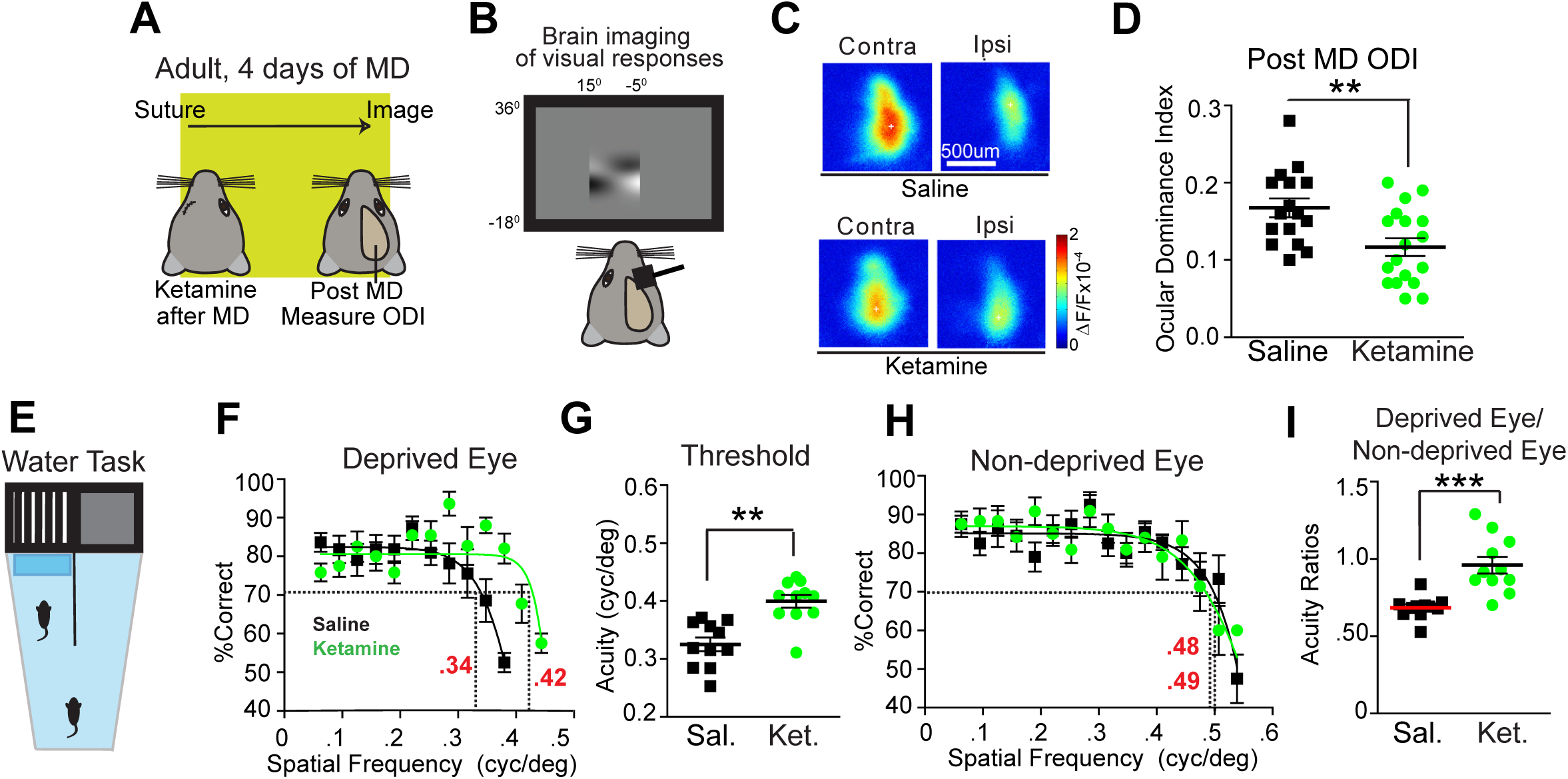
Ketamine reactivates adult ocular dominance plasticity and restores visual acuity in amblyopic animals. (A-D) Testing the effect of subanesthetic ketamine on ocular dominance plasticity during adulthood using intrinsic signal optical imaging. (A) Illustration of the experimental paradigm for assessing ocular dominance plasticity in adult (P90-120) mice, and the timeline of the experimental protocol. (A, left) Immediately after monocular deprivation (MD), saline or ketamine treatments are given, and 4 days later ocular dominance plasticity is measured after eyelid suture removal. (B) Intrinsic signal responses of visual cortex to contralateral versus ipsilateral eye stimulation are recorded and the ocular dominance index (ODI) is assessed during testing. The ODI is computed as (C-I)/(C+I) where C and I are the averaged map amplitudes calculated for contralateral and ipsilateral visual stimulation respectively. (C) Cortical response maps from mouse visual cortex. Representative contralateral and ipsilateral cortical response maps are shown from a control saline treated animal (top) and a ketamine treated animal (bottom) 4 days after treatment. In adult mice, which normally have relatively high responses to contralateral eye stimuli, 4 days of MD in saline treated mice does not induce changes in ODI, or a loss of contralateral eye input. (D) However, ODI is significantly reduced in ketamine treated adult animals (n=18, green) as compared to control saline treated animals (n=16, black). The data were pooled from cohorts of mice receiving either 10 or 50 mg/kg ketamine treatment, as they shared similar trends. (E-I) Testing the effect of ketamine on restoring functional visual perception in adult amblyopic mice. (E) Schematic of the visual water maze task. Mice have one eye sutured shut from P18-P32. At P32 the eye is re-opened and mice are treated with either saline or subanesthetic ketamine (10mg/kg; s.c.) every other day for three total treatments. At P90 mice begin water maze training, and are tested at P100. (F) Average performance on the visual task for all mice treated with either saline (black, n=11) or ketamine (10mg/kg; s.c., green, n=12) using the deprived eye. The performance curve is fit to a sigmoid. The x axis schematically represents the spatial frequency of the visual stimulus gratings. For saline treated mice the acuity threshold is 0.34 cpd. For ketamine treated mice the acuity threshold is 0.42 cpd. (G) Quantification of perceptual acuity across groups using acuity thresholds calculated from individual eyes. For deprived eyes ketamine treated animals (green, n=12) show significantly improved acuity thresholds compared to saline treated animals (black, n=11). (H) Average performance on the visual task for all mice treated with either saline (black, n=11) or ketamine (green, n=12) using the non-deprived eye, which reveals no significant differences across groups. For saline treated mice the acuity threshold is 0.49 cpd. For ketamine treated mice the acuity threshold is 0.48 cpd. (I) Acuity threshold ratios on the visual task for all mice treated with either saline (black, n=11) or ketamine (green, n=12). Acuity threshold ratios were determined by dividing the acuity threshold for the deprived eye by the threshold of the non-deprived eye. For mice treated with saline (black, n=11) or ketamine (green, n=12), the mean acuity thresholds ratio is significantly increased for ketamine-treated mice (*** p < 0.001; Student’s *t*-Test). Data represent mean ± SEM.

We tested whether ketamine treatment promotes visual functional recovery in adult amblyopic mice. Young mice were subjected to 2 weeks of monocular deprivation during the critical period (P18-32) to induce amblyopia (Davis et al., 2015). After eye sutures were removed, mice were then treated with saline or ketamine. At ∼3 months later, during adulthood (P90-P120), the acuity thresholds of individual deprived and non-deprived eyes were determined using the visual water task (Figure 1E-I) (Davis et al., 2015; Prusky et al., 2000). For the visual water maze task, mice were first trained to associate a hidden escape platform with a visual grating. Mice were then challenged during testing to find the platform as the spatial frequency of the grating was increased. Visual acuity of the deprived and non-deprived eye was assessed independently by covering one eye with an eye patch during testing. The perceptual threshold of vision in each eye for an individual mouse was then derived from the visual performances.

The average performance and threshold for vision through the deprived eye is dramatically improved with ketamine treatment (Figure 1F,G). This improvement is specific for the deprived eye as visual performance through the non-deprived eye is unchanged with ketamine treatment (Figure 1H). The control thresholds for vision in the non-deprived eyes are consistent with published data (Prusky et al., 2000). Acuity threshold improvements in the deprived eyes of ketamine treated mice are robust. Acuity threshold ratios for deprived/non-deprived eyes allow for an internal control within each animal (Figure 1I). These data support that subanesthetic ketamine treatment *in vivo* promotes recovery of visual acuity defects from amblyopia induced in the critical period of visual development. This has significant translational implications in developing new effective treatments for amblyopia.

### Cortical disinhibition evoked by subanesthetic in vivo ketamine

Changes in cortical inhibition exerted by inhibitory interneurons are essential for regulating cortical plasticity (Hensch, 2005; Kuhlman et al., 2013; Sun et al., 2016). We started by examining the time course of ketamine-evoked changes in cortical inhibition in adult mouse visual cortex. Ketamine undergoes metabolism *in vivo*, leading us to consider whether we would observe differences in ketamine responses after *in vivo* administration versus bath application. We first tested whether bath application of ketamine induces short-term changes in inhibitory input to excitatory pyramidal neurons in cortical layer 2/3 (L2/3) of the binocular region of mouse primary visual cortex, where the initial stage of critical period ocular dominance plasticity occurs in response to visual deprivation (Kuhlman et al., 2013). We performed whole-cell recording of electrically-evoked inhibitory postsynaptic currents (IPSCs) in L2/3 pyramidal neurons (Figure 2A), by preferentially activating L4->L2/3 feedforward projections to L2/3 inhibitory neurons through L4 electrical stimulation. We do not find any significant acute effects of bath applied ketamine on IPSCs (Figure 2B). We also tested the NMDAR antagonist MK-801 and do not observe any changes on IPSCs in pyramidal neurons after its bath application (Figure 2C). These results suggest that bath applied ketamine itself or the acutely action of blocking NMDARs does not modulate inhibitory synaptic inhibition to L2/3 pyramidal neurons in visual cortex.

**Figure 2.**
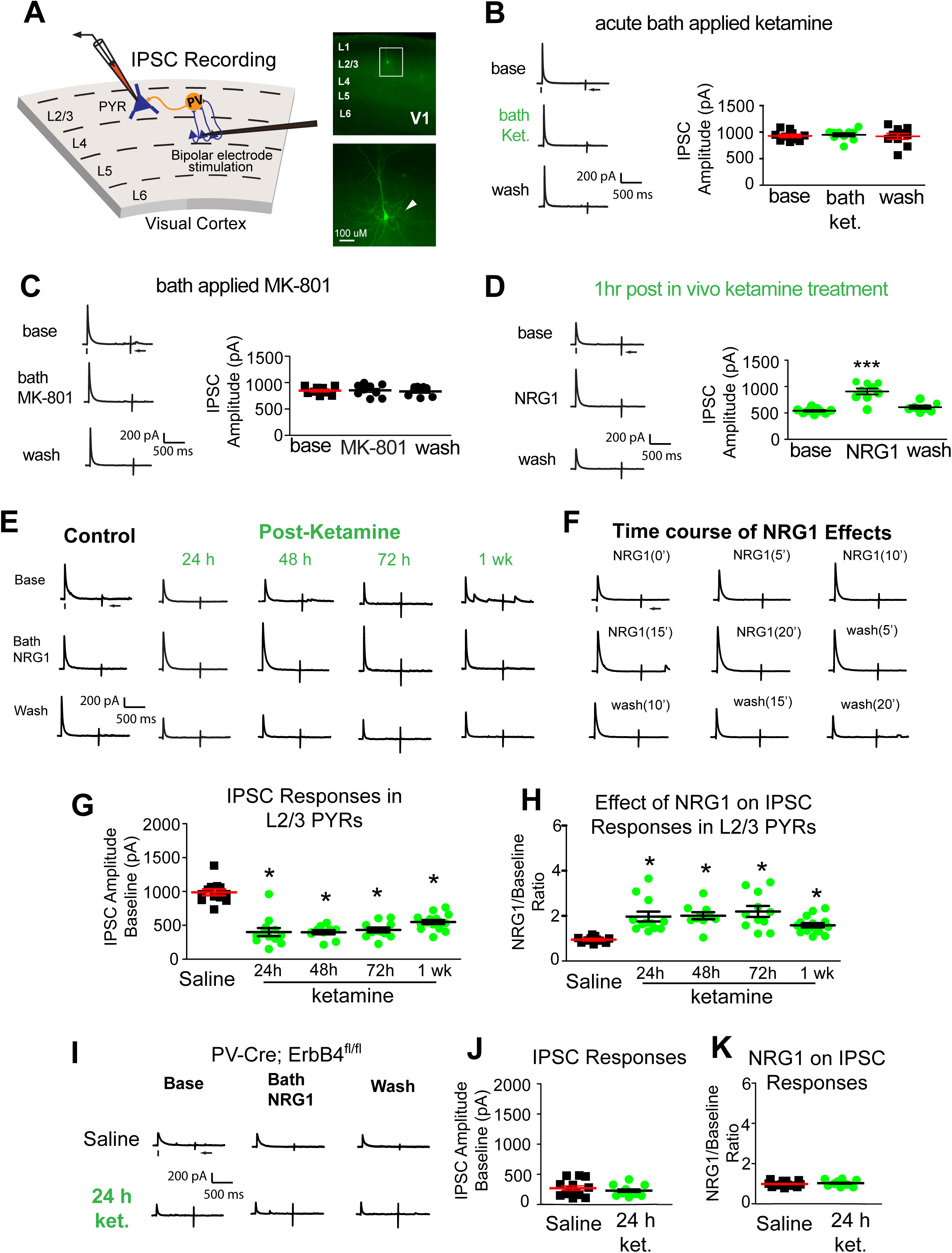
Ketamine evokes sustained cortical disinhibition, which is reversed with exogenous NRG1. (A) Schematic of recording inhibitory postsynaptic currents (IPSCs) in L2/3 pyramidal (PYR) neurons by preferentially activating L4→L2/3 feedforward projections to L2/3 PV neurons through L4 electrical stimulation. Recorded pyramidal neurons are filled with biocytin for post-hoc morphological confirmation. (B) Acute bath application of ketamine does not induce any change in inhibitory inputs to L2/3 pyramidal neurons (n=10 cells) in adult mouse visual cortex (P56 and above). (C) Similarly, acute bath application of MK-801 does not induce any change in inhibitory inputs to L2/3 pyramidal neurons (n=9 cells). (D) However, ketamine *in vivo* treatment (10 mg/kg; subcutaneous injection, s.c.) only 1 hour before recording reduces synaptic inhibition to L2/3 pyramidal cells. Bath application of recombinant NRG1 (5 nM) then reverses the effect of *in vivo* ketamine treatment and increased inhibitory inputs (n=9 cells). (E) Ketamine *in vivo* (10 mg/kg; s.c.) dramatically reduces evoked IPSCs to PYR neurons with sustained long-term effects at 24 hours (n=12 cells), 48 hours (n=10 cells), 72 hours (n=11 cells), and 1 week (n=12 cells) following treatment. In contrast to control excitatory neurons, ketamine evoked sustained decreases in IPSCs are acutely reversed with bath NRG1. (F) The time course of effects of acute application of NRG1. Bath application of recombinant NRG1 induces increases in inhibitory inputs to L2/3 pyramidal neurons with 10-20 minutes and this effect can be rapidly washed out within 20 minutes. (G,H) Summary data of average evoked IPSC amplitudes in L2/3 pyramidal neurons under the specified conditions (control saline, 24, 48 and 72 hours and 1 week after ketamine treatment). Acute NRG1 treatment increases average evoked IPSC amplitudes ratios in L2/3 pyramidal neurons under the specified conditions (control, 24, 48 and 72 hours and 1 week after ketamine treatment) after bath NRG1 application. (I-K) Genetic removal of NRG1 receptor ErbB4 in PV inhibitory cells prevents ketamine-induced decreases in evoked IPSCs in L2/3 excitatory neurons: neither ketamine injection (10mg/kg; s.c.) nor bath NRG1 application affects evoked IPSCs to L2/3 pyramidal neurons of PV-Cre; ErbB4^fl/fl^ mice. Representative IPSC traces shown in (I), and this data is summarized in (J, K). For each trial (A-L), electrical stimulation (1 ms, 20 uA) was applied, represented by a black tick beneath one example trace. For the example trace, the arrow indicates the current injection response to monitor access resistance during the experiment. (* p < 0.05, ** p < 0.01; Kruskal Wallis test, followed by Mann-Whitney U tests). Data represent mean ± SEM.

In contrast, we find rapid effects following *in vivo* subanesthetic ketamine treatment on decreasing inhibitory inputs to L2/3 excitatory cells. We injected mice with subanesthetic ketamine, and waited for 1 hour before making brain slices for IPSC recordings. We observe the evoked IPSC amplitudes are dramatically reduced to about 50% of control cells in non-treated cortex (Figure 2D). We next reasoned that the physiological impact of ketamine-mediated reduction in inhibitory input to excitatory pyramidal neurons might be reversed by enhancing NRG1/ErbB4 signaling. This was motivated by our previous findings that NRG1 strongly modulates evoked synaptic inhibition to pyramidal neurons in visual cortical slices when NRG1/ErbB4 signaling is downregulated (Grieco et al., 2019a; Sun et al., 2016). Exogenous recombinant NRG1 containing the epidermal growth factor (EGF) core domain of NRG1-β1 applied to the bath greatly reverses reductions in inhibitory inputs to the same group of L2/3 pyramidal cells recorded from visual cortex of the mice receiving *in vivo* ketamine treatment only shortly before (Figure 2D).

We then examined the sustained action of ketamine on the evoked inhibition of L2/3 pyramidal neurons at the different time points of 24, 48, 72 hours, and 1 week after single dose treatment. Inhibitory inputs to pyramidal neurons remain substantially reduced in ketamine treated mice at all the time points tested (Figure 2E,G). Ketamine evoked sustained decreases in evoked IPSCs are acutely reversible with bath NRG1 (Figure 2E,H); NRG1 increases and restores evoked IPSCs in pyramidal neurons in ketamine treated mice to control values. Bath acute NRG1 effects are robust and have a fast time course (Figure 2F). The response increases are clearly detectable within 10 minutes and reach a plateau around 20 minutes following NRG1 bath application. NRG1 potentiation persists with long duration in the continued presence of the peptide, but its effects are quickly washed out (Figure 2F). Note that exogenous NRG1 has no effects on IPSC amplitudes in normal control pyramidal neurons, suggesting that endogenous NRG1 signaling is sufficiently high to maintain synaptic inhibition to excitatory neurons in ketamine-untreated cortex.

The sustained disinhibitory action of ketamine on L2/3 pyramidal neurons, and the reversal of this effect by NRG1 may be specific to NRG1 signaling at ErbB4 receptors on PV neurons. To test the NRG1/ErbB4 signaling-mediated action of ketamine, we treated PV-Cre; ErbB4^fl/fl^ with ketamine, and then measured evoked IPSCs in visual cortical pyramidal neurons 24 hours after ketamine injection. The NRG1 cognate receptor ErbB4 is genetically removed from PV interneurons in PV-Cre; ErbB4^fl/fl^ mice (Long et al., 2003). While excitatory cells receive weak baseline inhibition, we find no significant differences in inhibitory input to L2/3 pyramidal cells after ketamine treatment in PV-Cre; ErbB4^fl/fl^ mice (Figure 2I,J). NRG1 does not alter evoked IPSCs in either ketamine treated or untreated PV-Cre; ErbB4^fl/fl^ mice (Figure 2I,K). These results support that ketamine-induced effects require PV specific ErbB4 for NRG1 signaling.

### NRG1/ErbB4 signaling in PV neurons reduced by ketamine

We asked if ketamine-evoked cortical disinhibition is mediated by downregulation of NRG1/ErbB4 signaling. To determine if a single subanesthetic ketamine treatment differentially modulates NRG1 or ErbB4 mRNA expression in PV inhibitory neurons and excitatory cells, we tested this using translating ribosome affinity purification (TRAP) to map cell type-specific mRNA expression changes (Zhou et al., 2013) (Figure 3A). NRG1 and ErbB4 mRNAs were measured from PV+ or Emx1+ neurons, which express the tagged ribosome from PV+ or non-GABA+ neurons, respectively (Figure 3B-C). Primary binocular visual cortex (bV1) was then harvested from the mice at 24, 48, 72 hours or 1 week after ketamine treatment. NRG1 expression in PV cells remains robust in control mouse cortex (Figure 3D); Emx1+ neurons express NRG1 mRNA at lower levels than PV interneurons (Figure 3F). In PV interneurons, ketamine induces a sustained down-regulation of NRG1 mRNA which returns to saline control baseline only after 1 week (Figure 3D). No changes are observed in Emx1+ neurons. The expression of ErbB4 mRNA does not change following ketamine treatment in either PV+ or Emx1+ neurons, albeit baseline ErbB4 is expressed at markedly higher levels in PV+ interneurons as compared to Emx1+ neurons (Figure 3E,G). The sustained and selective downregulation of NRG1 signaling in PV neurons after ketamine treatment strengthens its candidacy as the molecular basis of ketamine-mediated visual cortical plasticity during adulthood.

**Figure 3.**
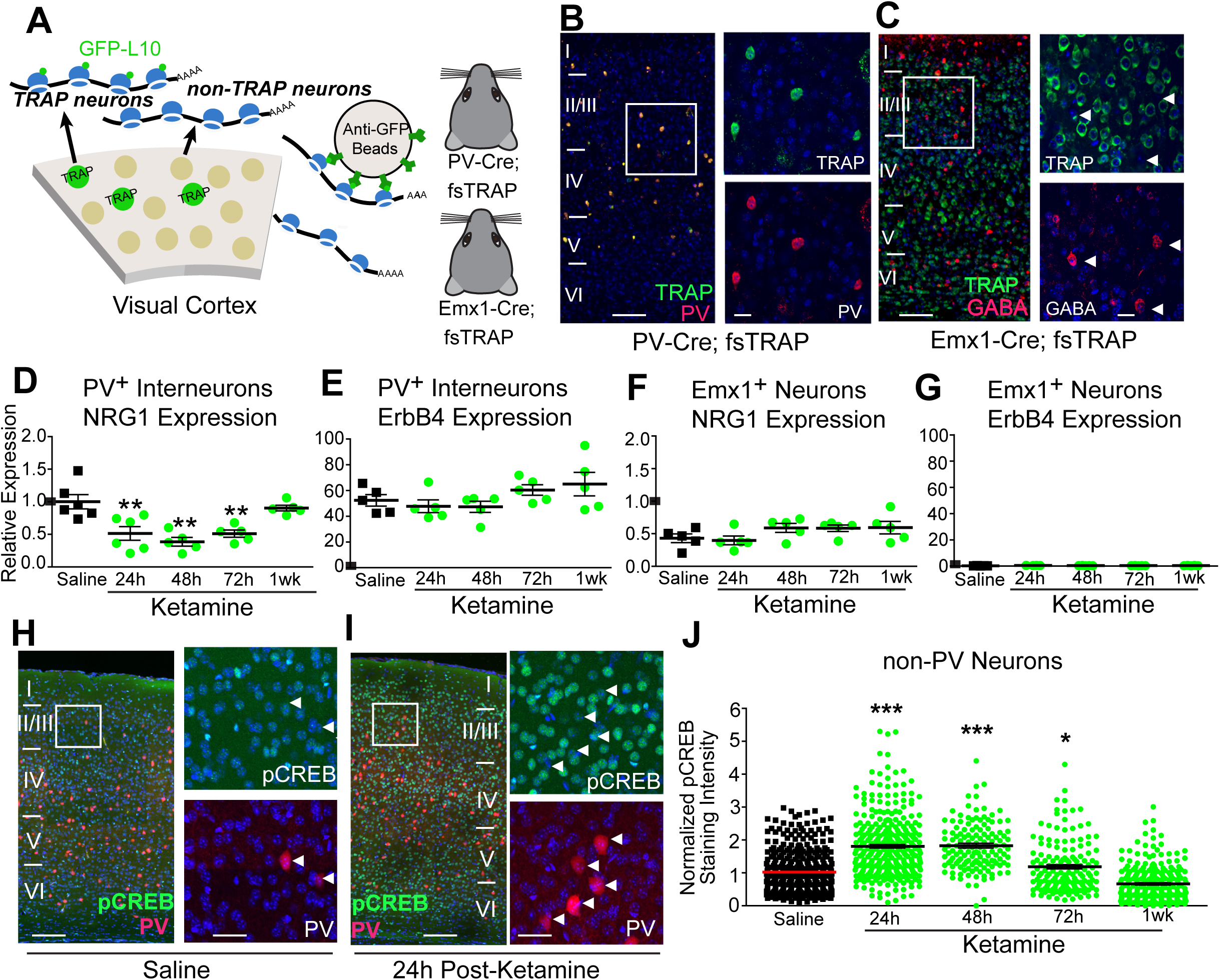
Ketamine induces sustained down-regulation of NRG1 in visual cortex PV inhibitory neurons. (A-G) Testing the effect of ketamine on cell-type specific NRG1 mRNA expression. (A) A schematic illustration of the translating ribosome affinity purification (TRAP) strategy. Using PV-Cre; fsTRAP mice, translating ribosomes (polysomes) from PV cells (green cells) have EGFP tags from the EGFP-L10 transgene. Lysis of all cells in the PV-Cre; fsTRAP cortex releases both tagged and non-tagged polysomes. The tagged polysomes are selectively captured on an anti-GFP affinity matrix and used for purification of PV-specific mRNAs associated with tagged polysomes. This strategy is also used to purify mRNA from Emx1+ neurons using Emx1-Cre; fsTRAP mice. Mice were treated with saline or ketamine (10 mg/kg; s.c.) at P56, then sacrificed 24, 48, 72 hours or 1 week later for fsTRAP extraction and qPCR analysis. (B) Confocal images of genetically labeled PV cells (green), PV immunolabeling (red) and their overlay in layer I-VI of mouse V1 in PV-Cre; fsTRAP-EGFP mice; scale bar = 200 µm. The white boxes indicate the region of V1 imaged at higher magnification (B, right); scale bar = 50 µm. (C) Confocal images of genetically labeled excitatory cells (green), GABA immunolabeling (red) and their overlay in layer I-VI of mouse V1 in Emx1-Cre; fsTRAP-EGFP mice; scale bar = 200 µm. The white boxes indicate the region of V1 imaged at higher magnification (C, right); scale bar = 50 µm. Arrowheads indicate GABA immunopositive cells. (D-G) All results were analyzed using the ddCt method with gapdh as an endogenous control, along with normalization to *nrg1* expression in P56 PV cells, as indicated by a horizontal mark on the y-axis, so that all panels (D-G) can be compared (Livak and Schmittgen, 2001). (D) Ketamine treatment significantly lowers NRG1 mRNA expression levels from PV neurons in visual cortex (n=5-6 samples; each sample is pooled from 5 mice for PV cell-specific data). (E) PV cell-specific ErbB4 mRNA expression levels do not significantly differ for mice treated with saline or ketamine. (F) Emx1+ cell-specific NRG1 mRNA expression levels from visual cortices (n=5 samples; each sample is pooled from 2 mice for Emx1+ cell-specific data) is not significantly changed with ketamine treatment. (G) Emx1+ cell-specific ErbB4 mRNA expression levels are not significantly changed with ketamine treatment. (H-J) Testing the effect of ketamine on pCREB expression in excitatory neurons. (H) Confocal images of genetically labeled PV cells (red), pCREB immunolabeling (green) and their overlay in layer I-VI of mouse V1 in P56 PV-Cre; Ai9 mice treated with saline; scale bar = 200 µm. The white box indicates the region of V1 digitally enlarged (H, right); Scale bar = 50 µm. White arrowheads indicate that PV neurons have very low pCREB immunoreactivity. (I) Confocal images of genetically labeled PV cells (red), pCREB immunolabeling (green) and their overlay in layer I-VI of mouse V1 in P56 PV-Cre; Ai9 mice 24 hours after ketamine treatment (10 mg/kg; s.c.); scale bar = 200 µm. The white boxes indicate the region of V1 digitally enlarged (I, right); scale bar = 50 µm. White arrowheads indicate that PV neurons have very low pCREB immunoreactivity. (J) Quantification of the increase in pCREB immunoreactivity of non-PV, putative excitatory neurons in visual cortex layer 2/3 at 24, 48 and 72 hours following ketamine treatment. After 1 week pCREB levels return to the saline control level. The overall normalized values of putative excitatory cells pooled from different mice were compared across treatment groups (n=8 for saline; n=3-7 for ketamine groups) (* p < 0.05, ** p < 0.01; Kruskal Wallis test, followed by Mann-Whitney U tests). Data represent mean ± SEM.

To determine if ketamine-mediated adult visual cortical plasticity and down-regulation of NRG1 expression in PV cells is associated with increases in molecular correlates of enhanced neural excitability via cortical disinhibition, the levels of phosphorylated cAMP response element transcription factor (pCREB) were measured by immunostaining after ketamine treatment (Figure 3H-J). In excitatory cortical neurons, the regulation of expression of specific genes depends on activation of CREB by phosphorylation at Ser133 (pCREB); CREB phosphorylation at Ser133 occurs in response to increased neural activity (Cohen and Greenberg, 2008). While pCREB levels in PV neurons are very low, a single subanesthetic ketamine dose increases pCREB levels in putative excitatory neurons (non-PV cells) in visual cortex (Figure 3H,I). We quantified and compared pCREB immunostaining fluorescence intensity from layer 2/3 excitatory neurons in saline, and ketamine-treated mice. We find that compared to control measurements from non-ketamine treated mice, an increase of pCREB activity levels in excitatory neurons is sustained for at least 72 hours after ketamine treatment. pCREB levels return to saline-control baseline levels at 1 week following ketamine treatment (Figure 3J). These results indicate that increased excitatory neuron activity via cortical disinhibition is sustained in visual cortex following a subanesthetic ketamine treatment, and provide further motivation to test the mechanisms underlying ketamine-mediated cortical disinhibition.

### Sustained loss of excitatory inputs to PV neurons by ketamine

To determine if PV cell excitatory input circuitry is the synaptic locus leading to cortical disinhibition, we tested the effects of ketamine and NRG1 treatment on excitatory inputs to PV cells. We measured the strength of excitatory inputs to local L2/3 PV neurons using laser scanning photostimulation (LSPS) in brain slices of adult binocular visual cortex (Kuhlman et al., 2013; Xu et al., 2016) (Figure 4A-D). The LSPS approach is effective for detailed local circuit mapping. It involves first recording from a single neuron, then sequentially stimulating at surrounding sites to evoke action potentials from neurons in those sites through spatially restricted UV laser-evoked glutamate release; recording from the potential postsynaptic neuron allows us to determine whether there are synaptic inputs from those stimulation sites (Figure S2). LSPS experiments were performed in visual cortical slices prepared from control mice and *in vivo* ketamine treated adult mice at the different time points of 24, 48, 72 hours, and 1 week after treatment. We find that control PV neurons receive robust excitatory inputs from L4 and upper L5, as well as from L2/3 (Figure 4E,F). In ketamine treated mice, however, the excitatory drive to PV neurons is dramatically reduced 24, 48 and 72 hours later (Figure 4E, top row). Excitatory input onto PV neurons tends to recover partially at 1 week after ketamine treatment (Figure 4E, F). These results demonstrate that excitatory input to PV neurons is attenuated following *in vivo* ketamine treatment and that this response is sustained for around 1 week.

**Figure 4.**
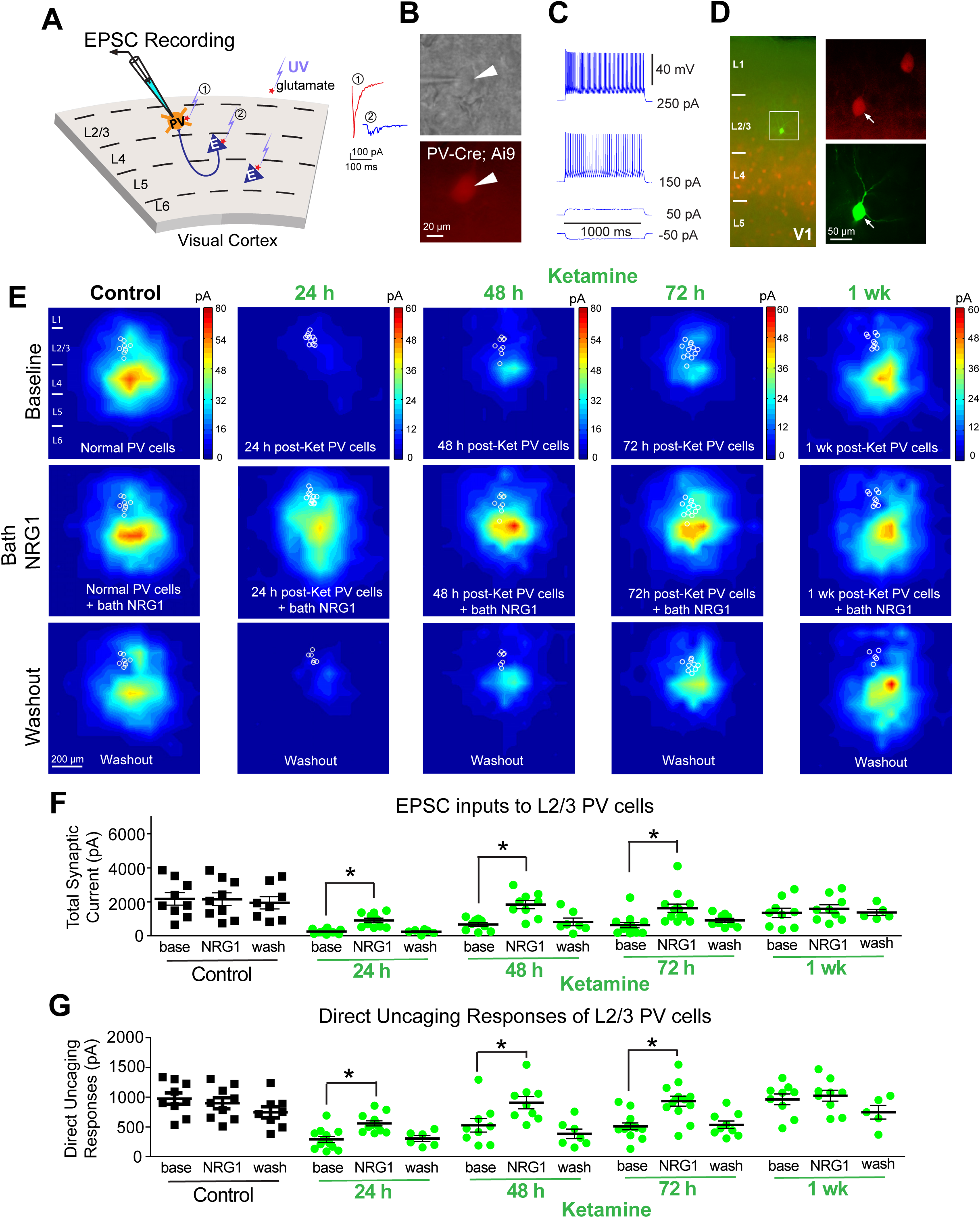
Ketamine evokes sustained loss of excitatory inputs to PV interneurons, which is restored with NRG1. (A) Schematic of laser-scannning photostimulation (LSPS) mapping of cortical synaptic connections to individually recorded PV neurons in V1 slices. LSPS maps the broad spatial pattern of synaptic inputs to the neuron of interest, and distinguishes direct uncaging responses (1, red) to assess glutamate-mediated excitability/responsiveness at perisomatic locations, and synaptically mediated EPSC responses (2, cyan) to assess synaptic inputs from presynaptic neuronal spiking. Cortical layers of 1, 2/3, 4, 5, and 6 in the brain slice are indicated as L1, L2/3, L4, L5, and L6. (B) Targeted recordings of PV neurons are facilitated by tdTomato expression in PV-Cre; Ai9 mouse slices. (C) PV neurons are further identified by their fast-spiking profile. (D) In addition, recorded neurons are injected with biocytin for further post-hoc morphological analysis. (E) Group-averaged, excitatory input maps of PV cells recorded at the specified conditions. White circles in each map represent individual PV neurons. The color scales code integrated excitatory input strength (blue = low, red = high) and applies to all other maps in the same condition. (E, left column) An acute bath application of NRG1 (5 nM) does not significantly modulate local excitatory synaptic inputs onto PV neurons in V1 of saline-treatment control mice. Averaged excitatory input maps of L2/3 PV cells are shown before (top), during bath NRG1 (20 min after NRG1 application) (middle), and after washout of bath NRG1 (bottom). (E, middle columns) Reduced excitatory inputs to PV neurons are seen at 24, 48 and 72 hours after ketamine treatment (10 mg/kg; s.c.), but not after 1 week. (E, middle row) Excitatory inputs to PV neurons in ketamine treated mice are restored by acute bath application of NRG1. (E, bottom row) This acute restoration by NRG1 is eliminated by washout of the bath NRG1. (F) Summary data of average total synaptic input strength measured for L2/3 PV neurons under the specified conditions (control, 24, 48 and 72 hours and 1 week after ketamine treatment). (G) Direct uncaging responses before and during bath NRG1 for PV neurons in control versus ketamine treated mice. Peak direct responses were measured, which are not affected by overriding synaptic inputs (* p < 0.05; Kruskal Wallis test, followed by Mann-Whitney U tests). Data represent mean ± SEM.

We then examined whether enhanced NRG1 signaling restores excitatory input onto these PV cells. Bath applied NRG1 does not alter resting membrane potential or intrinsic membrane excitability in PV cells under control conditions or following ketamine treatment (Figure S3). After baseline recording, we applied NRG1 in the bath to cortical slices for 20 minutes. For control PV cells in mice that were not treated with ketamine, neither local excitatory synaptic input currents nor direct uncaging response (as measured by responses to glutamate uncaging at perisomatic regions) of PV neurons are significantly altered with the NRG1 application (Figure 4E,F). In contrast, both excitatory synaptic input currents and direct uncaging responses of PV neurons are strongly enhanced by bath NRG1 in ketamine-treated mice at 24, 48 and 72 hours after ketamine injection (Figure 4E-G). As fast excitatory synaptic currents are mediated by postsynaptic AMPA receptors in PV neurons (Figure S2), this result also indicates that downstream NRG1 signaling modulates synaptic response through AMPA receptors.

To confirm the specificity of the action of ketamine on NRG1/ErbB4 signaling in PV neurons, we also mapped the strength of presynaptic excitatory inputs to L2/3 excitatory neurons in binocular visual cortex after *in vivo* ketamine treatment. In contrast with the modulation of excitatory inputs to PV cells, ketamine treatment or acute application of recombinant NRG1 does not significantly modulates local excitatory synaptic inputs or direct uncaging responses of L2/3 excitatory pyramidal neurons (Figure S4). Neither ketamine treatment nor exogenous NRG1 application alters resting membrane potential or intrinsic membrane excitability of L2/3 pyramidal neurons (Figure S4). These results demonstrate that excitatory inputs to L2/3 pyramidal neurons are not altered following *in vivo* ketamine treatment. Together, combined with our finding of ketamine-evoked cell-type specific NRG1 downregulation, our data support that the sustained effect of ketamine is specifically mediated through NRG1/ErbB4 signaling in PV inhibitory neurons.

### Effects of the ketamine metabolite HNK

Given our contrasting effects of bath-applied ketamine and *in vivo* ketamine treatment, we consider the recent finding that the metabolism of ketamine to (2R, 6R)-HNK is essential for its antidepressant effects (Zanos et al., 2016). We tested the hypothesis that the HNK is the biologically active metabolite of ketamine by determining whether HNK treatment effects are similar to ketamine. Acute bath application of (2R, 6R)-HNK (100 µM) did not induce significant changes in evoked IPSC strength to L2/3 excitatory pyramidal neurons (Figure 5A, B), which is consistent with the previous finding that it takes about an hour for the HNK to increase AMPA receptor-mediated excitatory post-synaptic potentials recorded from the CA1 region of hippocampal slices (Zanos et al., 2016). Similar to *in vivo* ketamine treatment, we find that inhibitory inputs to L2/3 pyramidal neurons are greatly reduced in visual cortex in HNK treated mice 24 hours following treatment (Figure 5A,C). Bath application of recombinant NRG1 significantly restores those inhibitory inputs, as measured by whole-cell recording of electrically-evoked IPSCs in L2/3 pyramidal neurons in visual cortex of HNK treated mice (Figure 5A,C). To further compare HNK and ketamine responses, we quantified excitatory inputs to L2/3 PV interneurons in the visual cortex using LSPS. We find that excitatory inputs onto PV neurons are largely lost 24 hours following HNK treatment (Figure 5D-G). Counteracting HNK effects, bath application of NRG1 restores the excitatory drive onto PV neurons rapidly (Figure 5E-G).

**Figure 5.**
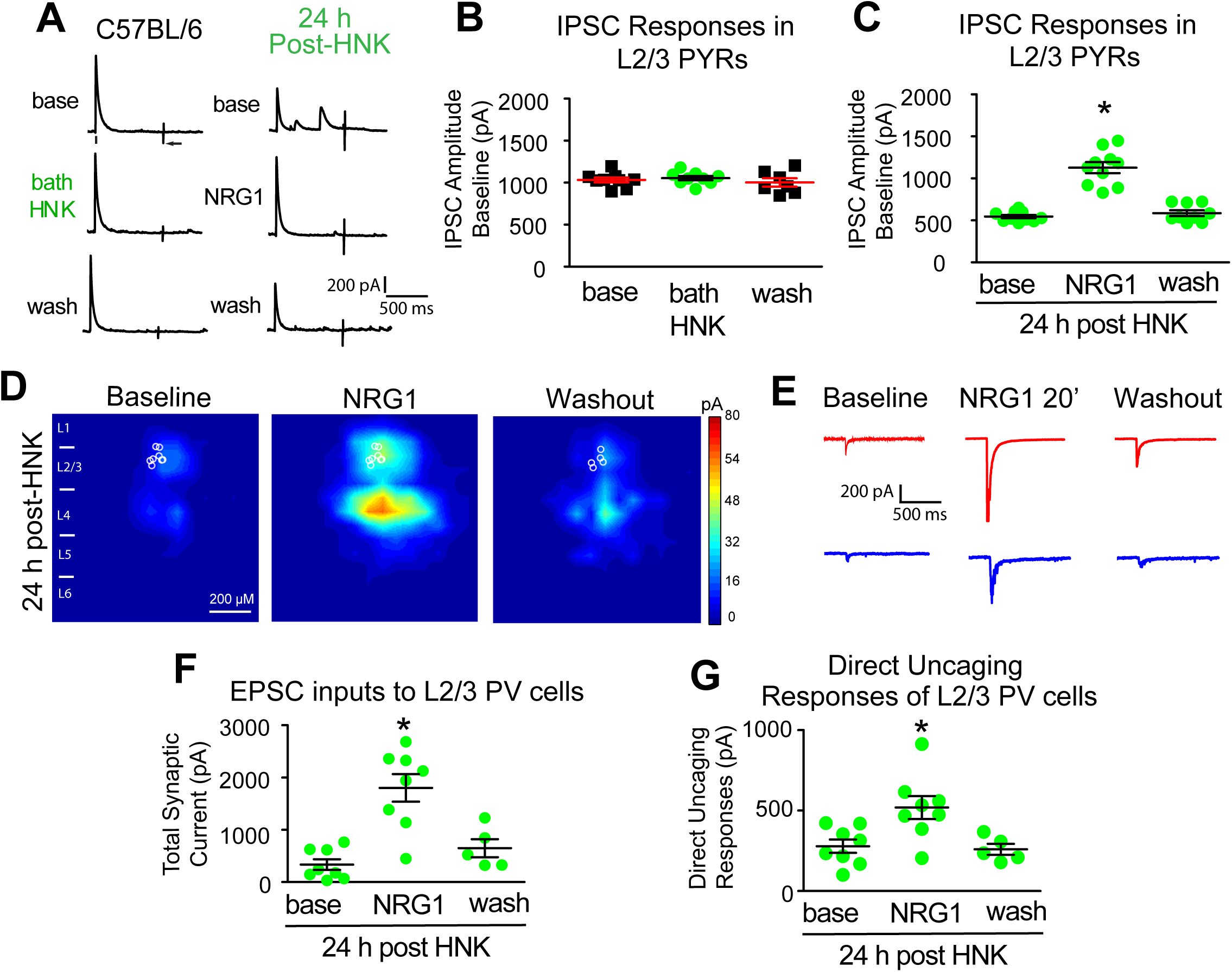
HNK evokes sustained reduction of cortical inhibition and loss of excitatory inputs to PV inhibitory neurons, both of which are restored with NRG1. (A) Acute bath application of (2R, 6R)-HNK (100 μM) does not induce significant changes in evoked IPSC strength to L2/3 excitatory pyramidal neurons. IPSCs were recorded in L2/3 pyramidal (PYR) neurons through L4 electrical stimulation as shown in Figure 2A. HNK treatment (10 mg/kg; s.c.) at 24 hours before recording induces a reduction of IPSCs to L2/3 pyramidal neurons. Acute bath application of NRG1 (5 nM) reversed the reduced inhibition induced by HNK treatment. (D) LSPS mapping of local excitatory synaptic inputs to individually recorded PV neurons. (E) Illustration of example responses of a PV neuron in HNK treated mouse cortex with LSPS in a perisomatic location (red traces) and in a presynaptic input site (blue traces) at baseline, NRG1 bath application for 20 minutes and washout. (F-G) Both total synaptic inputs and direct uncaging response are reduced 24 hours after HNK treatment *in vivo*. Acute bath application of NRG1 (5 nM) significantly restore excitatory synaptic inputs onto PV neurons in V1 of HNK treated mice (* p < 0.05; Kruskal Wallis test, followed by Mann-Whitney U tests). Data represent mean ± SEM.

### Longitudinal examination of ketamine effects on population neurons

Extending our slice mapping results to *in vivo* physiology, we performed 2-photon calcium imaging of population neurons in mouse binocular visual cortex. We recorded from mice that express GCaMP6s in excitatory neurons (Wekselblatt et al., 2016). This allows us to probe the GCaMP6-based neural activity of population single excitatory neurons (Figure 6A). We longitudinally tracked and measured the effects of ketamine on excitatory neuron disinhibition *in vivo* in the visual cortex of awake head-fixed mice. We first performed baseline recordings to ensure the stability of responses (Figure S5), then treated mice with ketamine and collected data at different time points after treatment (Figure 6B). The *in vivo* activity of excitatory neurons measured by calcium event frequency significantly increases at 24, 48, and 72 hours following ketamine treatment (Figure 6C,D,G). At 1 week following a single ketamine treatment, excitatory neuron responses *in vivo* return to baseline levels (Figure 6C,D,G). *In vivo* 2-photon recordings from excitatory neurons show that ketamine treatment induces sustained disinhibition of population excitatory neuron activity at single cell resolution. This is supported by visually-evoked potential (VEP) recordings (Kaplan et al., 2016) from binocular visual cortex in un-anaesthetized and awake mice, where the sustained effects of ketamine on cortical disinhibition *in vivo* were also measured (Figure S6). Ketamine treatment significantly increases the VEP amplitudes 24 and 48 h after treatment. Furthermore, *in vivo* exogenous NRG1 treatment before ketamine injection prevents ketamine-induced increases in VEP amplitudes (Figure S6), which suggests exogenous NRG1 acutely blocks ketamine-induced disinhibition through enhancement of cortical inhibition (Sun et al., 2016).

**Figure 6.**
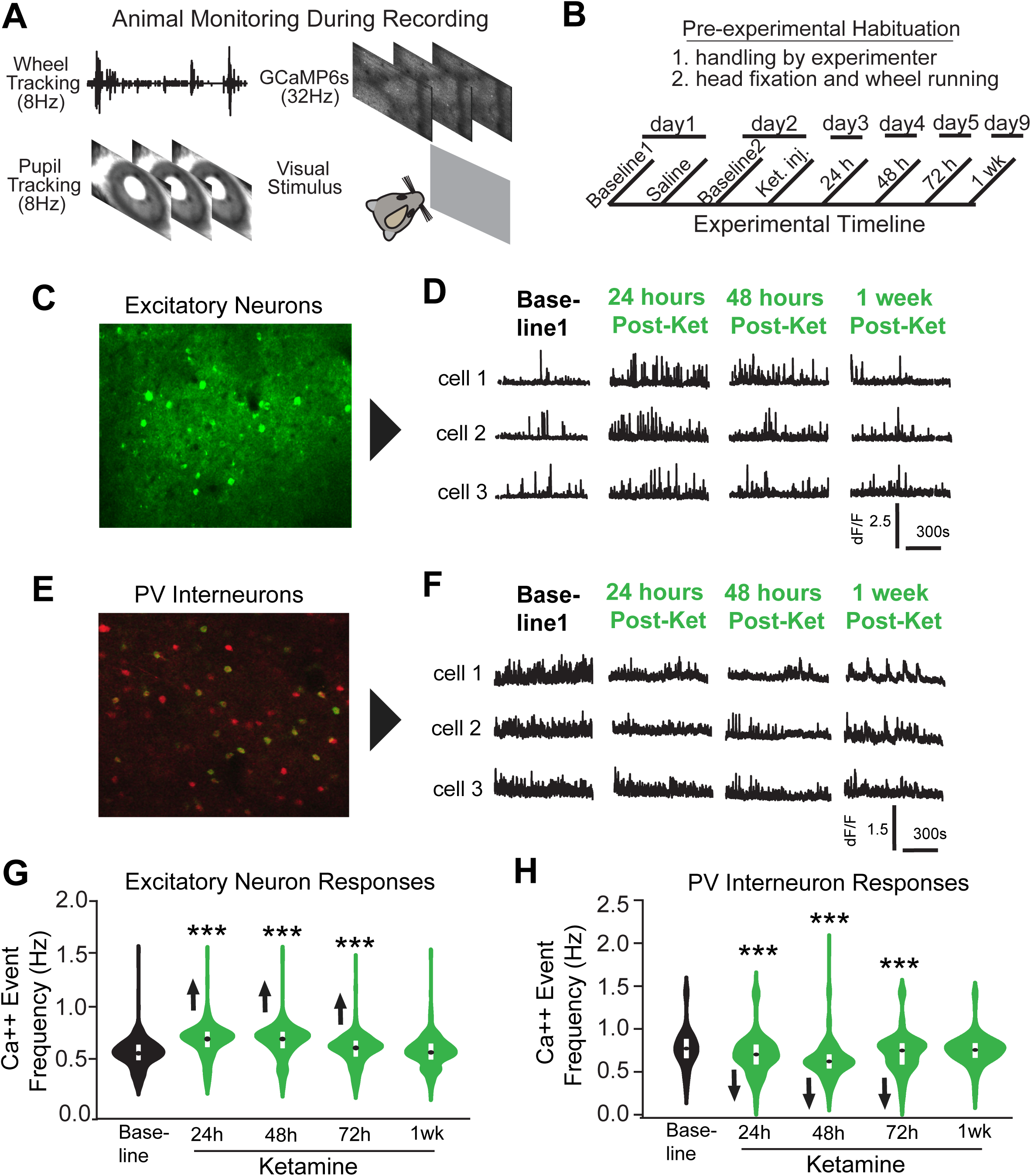
*In vivo* population calcium imaging reveals sustained disinhibition of excitatory neurons and inhibition of PV interneurons by subanesthetic ketamine in the visual cortex of awake head-fixed mice. (A) During recording sessions, wheel tracking was collected to determine animal running behavior, GCaMp6s calcium signals were recorded from excitatory neurons or PV interneurons using 2-photon calcium imaging, pupil tracking was also performed to monitor pupil sizes and eye movements, and data about the visual stimulus was recorded to ensure time stamping accuracy. (B) Illustration of the timeline of the experimental protocol. Briefly, baseline neuron activity is recorded and then ketamine is administered (10mg/kg; s.c.) to adult mice expressing GCaMP6s, and subsequent recordings are made at 24, 48, 72 hours and 1 week following treatment. (C) A representative maximum intensity Z-projection image of a swath of visual cortex from a GCaMP6s transgenic mouse expressing GCaMP6s in excitatory neurons (Wekselblatt et al., 2016). (D) Example calcium transient event data of three representative bV1 excitatory neurons from the experiment described in B. (E) A representative maximum intensity Z-projection image of a swath of visual cortex from a PV-Cre; Ai163 mouse (Daigle et al., 2018) expressing GCaMP6s (green) in PV cells (red). (F) Example calcium transient event data of three representative bV1 PV neurons from the experiment described in B. (G) Average calcium event frequencies of excitatory neurons across several recording sessions either before (baseline) or after ketamine treatment (24, 48, 72 hours and 1 week). (H) Average calcium event frequencies of PV inhibitory neurons across several recording sessions either before (baseline) or after ketamine treatment (24, 48, 72 hours and 1 week) (* p < 0.05; Kruskal Wallis test, followed by Mann-Whitney U tests). Data represent mean ± SEM.

We verified that ketamine enhances cortical excitability *in vivo* by inhibiting neural activity of PV inhibitory neurons by performing 2-photon longitudinal recording. Calcium activity of PV cells was measured using the genetically encoded calcium indicator GCaMP6s selectively expressed in PV neurons (Daigle et al., 2018). We determined that PV activity was stable (Figure S5), and then treated mice with ketamine and collected data at different time points after treatment (Figure 6B). The *in vivo* activity of PV neurons measured by calcium event frequency is significantly reduced at 24, 48, and 72 hours following ketamine treatment (Figure 6E,F,H). At 1 week following ketamine treatment, PV neuron responses return to baseline levels (Figure 6E,F,H). These data indicate that ketamine-mediated disinhibition of visual cortex is induced by reducing the population activity of PV interneurons.

Ketamine-evoked NRG1 signaling downregulation in PV neurons and ketamine associated disinhibitory effects *in vivo* are further supported by our *in vivo* miniscope imaging results in awake freely moving PV-Cre; ErbB4^fl/fl^ mice. We imaged GCaMP6-based neural activity of excitatory neurons in wild type and PV-Cre; ErbB4^fl/fl^ mouse visual cortex using head-mounted miniscopes (Cai et al., 2016; Ghosh et al., 2011; Sun et al., 2019) (Figure 7A-C; Figure S7). To determine the effects of *in vivo* manipulation of NRG1/ErbB4 signaling in PV cells on excitatory cortical activity, we selectively expressed GCaMP6 in excitatory neurons with AAV-CaMKII2a-GCaMP6f into mouse visual cortex. During experimentation, mice freely explored a small open arena. No differences in behavior are observed for PV-Cre; ErbB4^fl/fl^ mice and control mice (Figure 7D,E,H). Robust and significant increases in the amplitudes of excitatory neuron responses are found in PV-Cre; ErbB4^fl/fl^ mice as compared to controls (Figure 7F,G,I,J). Thus, combined with circuit mapping data, our *in vivo* imaging results support that ketamine acts on PV cell-specific NRG1/ErbB4 signaling in the regulation of neural circuit inhibition in adult visual cortex.

**Figure 7.**
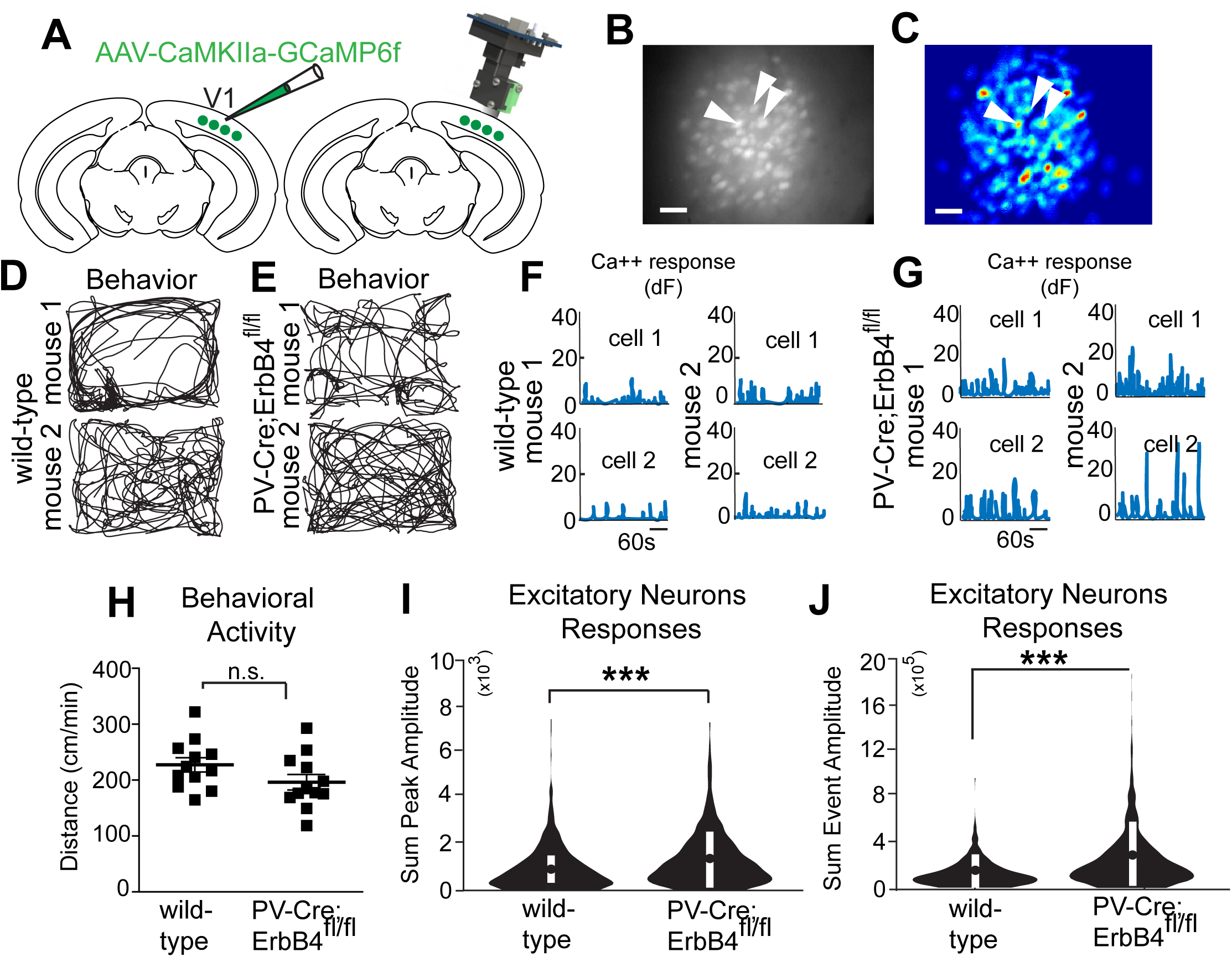
NRG1/ErbB4 signaling in PV interneurons is implicated in regulation of cortical inhibition. (A) A schematic illustration of AAV1 injection for targeted expression of GCaMP6f in bV1 excitatory neurons. A schematic illustration of a miniaturized fluorescent microscope (miniscope) used to image in vivo calcium signals in bV1 excitatory neurons in awake and freely behaving mice. The GRIN lens, implanted over bV1 at 2 weeks after an AAV1-Camk2a-GCaMP6f injection into bV1, is shown in grey. With the miniscope being fixed in place with a baseplate, reliable imaging of the same group of neurons over days can be repeatedly performed. Recording of neural and behavioral activity was performed during 5-10 minute sessions. (B) Representative maximum intensity Z-projection image showing recorded bV1 neurons from a combined recording session, scale bar = 50μm. (C) A spatial profile image shows extracted neurons and their contours (see white arrows) using the CNMF-E algorithm (see Materials and Methods for details) based on the concatenated video in (B), scale bar = 50μM. (D) Behavioral tracking data (black lines) of two representative wild type mice. The mice were introduced to a 35cm x 25cm behavioral arena under low lighting conditions (<20 lux), and miniscope and behavioral recordings were made. (E) Behavioral tracking data of two representative PV-Cre; ErbB4^fl/fl^ mice. (F) Calcium transient event data of two representative bV1 neurons from 2 different wild type mice (top, neuron 1; bottom, neuron 2). (G) Calcium transient event data of two neurons from two PV-Cre; ErbB4^fl/fl^ mice (top, neuron 1; bottom, neuron 2). (H) There is no significant difference in behavior (normalized distance; cm/min) between wild type and PV-Cre; ErbB4^fl/fl^ mice. (I) Average peak calcium event amplitudes, as determined by event amplitudes (Figure S7; see Methods for details), for neurons from wild type mice (n = 360 cells) as compared to PV-Cre; ErbB4^fl/fl^ mice (n = 627 cells). (J) Average summed event amplitudes for neurons from wild type mice (n=260) as compared to PV-Cre; ErbB4^fl/fl^ mice (n=627) (*** p < 0.001; Student t tests). Data represent mean ± SEM.

## Discussion

Subanesthetic *in vivo* ketamine reactivates adult visual cortical plasticity with features similar to the early postnatal critical period. Furthermore, subanesthetic ketamine treatment promotes adult functional recovery of visual acuity defects from amblyopia induced during the critical period of visual development. Re-opening adult cortical plasticity by subanesthetic ketamine treatment depends on a novel and critical role for NRG1/ErbB4 signaling at molecular, cellular and circuit levels.

Consistent with the notion that ketamine actions are mediated by neural activity-dependent molecular signals, the effect of a single ketamine treatment is rapid and effective. This is in contrast to chronic fluoxetine administration that requires daily dosing and several weeks of treatment (Maya Vetencourt et al., 2008). We also find that the ketamine metabolite, (2R, 6R)-HNK has rapid and long-lasting effects on synaptic plasticity, mimicking ketamine treatment. These pre-clinical findings provide motivation for testing subanesthetic ketamine and its metabolites for treating adult amblyopia. In particular, HNK is of strong interest as a therapeutic agent as it lacks the abuse liability and the dissociative side effects of ketamine.

Ketamine-induced effects in adult cortex strongly resemble features of monocular deprivation in the cortex of critical period animals. Previously, we showed that activity-dependent NRG1 molecular signaling and sensory experience interact to rapidly shape functional circuit connections in the visual cortex during the developmental critical period (Sun et al., 2016). We now find that subanesthetic *in vivo* ketamine treatment evokes a rapid reduction (as short as 1 hour) in synaptic inhibition to excitatory neurons in visual cortical layer 2/3. A prominent theory of the mechanism of action of ketamine argues that ketamine acts via direct inhibition of NMDARs localized to interneurons. Our data collected in visual cortex indicate that the neural mechanism of cortical disinhibition is independent of acute NMDAR antagonism as bath-applied ketamine itself or MK-801 does not modulate synaptic inhibition to L2/3 pyramidal neurons in visual cortex. A dramatic reduction (50%) of evoked IPSC inputs to excitatory neurons occurs within 24 hours of post-ketamine treatment or monocular deprivation. Ketamine induces increases in pCREB in excitatory neurons, indicating increased neural activity. The initial reduction of cortical inhibition alters visual cortical excitation / inhibition balance and establishes the conditions necessary for the experience-dependent ocular dominance plasticity.

We show that ketamine treatment downregulates neuregulin-1(NRG1)/ErbB4 signaling in PV neurons, which correlates with the rapid and sustained decreases in synaptic inhibition to excitatory neurons for up to a week. In contrast, ketamine has no effect on NRG1 expression in excitatory neurons. The acute and selective downregulation of NRG1 signaling in PV neurons induced by ketamine supports that this pathway serves as the molecular basis of the rapid retraction of excitatory inputs to PV cells and their reduced activity. Our parallel circuit mapping experiments identify the synaptic and circuit mechanism through which ketamine downregulates NRG1 signaling to reduce AMPA receptor-mediated synaptic inputs to PV neurons and PV cell-mediated inhibition to excitatory cells. This is supported by our findings that exogenous NRG1 rapidly restores normal excitatory input onto PV cells and enhances cortical inhibition in ketamine-treated cortex. In accordance with our previous study (Sun et al., 2016), rapid NRG1 potentiation of glutamate-evoked responses of PV cells in ketamine-treated cortex is consistent with the rapid insertion of the intracellular pool of AMPA receptors and increased clustering of AMPA receptors at the membrane surface of PV neurons. Furthermore, ketamine-induced effects require ErbB4 expression in PV neurons. Synaptic inhibition to cortical excitatory neurons is not modulated by NRG1 in either ketamine treated or untreated PV ErbB4 null mouse cortex. These results support the conclusion that ketamine-induced effects require PV specific ErbB4 for NRG1 signaling. The synaptic and circuit mechanisms through which ketamine modulates cortical inhibition through NRG1 signaling mechanisms may be generalized across cortical regions. Our findings offer mechanistic insights for previous studies in the hippocampus showing that within approximately an hour, (2R, 6R)-HNK induces a robust increase in AMPAR-mediated excitatory post-synaptic potentials (EPSPs) recorded from the CA1 region of hippocampal slices, which is sustained after washout of the drug (Zanos et al., 2016). The acute and sustained cortical disinhibition via downregulated NRG1 signaling may account for NMDAR inhibition-independent antidepressant actions of ketamine metabolites.

Our novel findings focus on PV-inhibitory neurons as the initial functional locus for the molecular and circuit mechanisms underlying ketamine’s effects. The PV-cell specific NRG1/ErbB4 mechanism identified in our work unifies previous proposals for molecular signaling mechanisms of ketamine’s action. The cortical disinhibition evoked by PV-specific NRG1 downregulation increases pCREB activity in excitatory neurons, which promotes BDNF production (Cohen and Greenberg, 2008). Ketamine-evoked changes in PV NRG1/ErbB4 signaling may initiate BDNF signal changes in excitatory neurons to mediate ketamine’s antidepressant responses. The BDNF receptor TrkB can activate further downstream pathways, including the serine-threonine kinase mTOR, which is also involved in neural plasticity (Duman et al., 2012). Activation of mTOR in excitatory cells is involved in promoting ketamine-induced dendrite spine growth (Li et al., 2010; Moda-Sava et al., 2019). mTOR signaling inhibits GSK3, which is the target of many mood stabilizers and antidepressants (Beurel et al., 2015). Intriguingly, similar to fluoxetine, lithium treatment of bipolar disorder requires at least several weeks of chronic administration and ultimately modulates GSK3 (Jope, 2003). While our data support that ketamine-evoked acute and sustained cortical disinhibition is independent of blockade of NMDA receptors in the visual cortex, it is relevant that ketamine can inactivate the calcium dependent eEF2 kinase to derepress BDNF synthesis and increase neural plasticity (Autry et al 2011). We identify here a PV-inhibitory neuron-directed molecular and circuit mechanism for ketamine’s effects on cortical plasticity that may also be informative for treating depression and affective disorders.

While this study has a justified focus on PV inhibitory neurons, it will also be important to examine whether and how ketamine modulates synaptic plasticity and function of non-PV inhibitory neuron types. Considering PV neurons can be inhibited by somatostatin-expressing interneurons and VIP interneurons preferentially inhibit somatostatin-expressing cells (Kepecs and Fishell, 2014), further studies are required for expand our understanding of ketamine induced effects beyond PV interneurons and excitatory neurons.

In summary, ketamine treatment promotes adult cortical plasticity and functional recovery from amblyopia by way of downregulation of PV-specific NRG1 signaling, which results in PV excitatory input loss and cortical disinhibition. Fast and sustained ketamine actions show promise for therapeutic applications that rely on re-activating adult cortical plasticity.

### Experimental Procedures

#### Animals

All experimental procedures and protocols were approved by the Institutional Animal Care and Use Committee of the University of California, Irvine. To enable PV specific labeling and mRNA expression analysis, PV-IRES-Cre knock-in (Jackson Laboratory, stock #008069) mice were crossed to fsTRAP mice (Zhou et al., 2013)(Jackson Laboratory, stock #022367) to generate PV-Cre+/-; fsTRAP mice, in which translating polyribosomes of PV cells are tagged with EGFP from the GFP-L10 transgene. To enable excitatory neuron labeling and mRNA expression analysis, Emx1-Cre mice were crossed to fsTRAP mice to generate Emx1-Cre+/-; fsTRAP mice. Emx1 is predominantly expressed in cortical excitatory neurons(Gorski et al., 2002). For all experiments, mice were hemizygous for both transgenes. To genetically label PV cells PV-Cre mice were crossed with Ai9 tdTomato reporter knock-in mice (Jackson Laboratory, stock #007905). To generate ErbB4 conditional knockout mice, mice homozygous for loxP-flanked alleles (Long et al., 2003) were crossed with PV-Cre mice to produce PV-Cre+/-; ErbB4flx/+ mice. These mice were then crossed back to the homozygous loxP-flanked ErbB4 mice to produce PV-Cre+/-; ErbB4 flx/flx mice. To selectively express GCaMP6s in PV cells, PV-Cre mice were crossed to Ai163 mice to generate PV-Cre; Ai163 mice. The animals (2-5 mice per cage) were housed in a vivarium room with a 12-h light/dark cycle with access to food and water ad libitum. Please see Supplementary Table 1 for detailed experimental and animal information. The mice were randomly assigned to groups with treatment of either saline or subanesthetic ketamine. Unless specified otherwise in some Figure 1 related experiments, a single treatment of ketamine (10 mg/kg; ketamine hydrochloride; VedCo., Inc.) was used for all the experiments. The (2R, 6R)-HNK dosage (10 mg/kg; Tocris Bioscience) of the published study (Zanos et al., 2016) was used for our *in vivo* HNK treatment. During some of the physiological experiments, we performed *in vivo* exogenous NRG1 treatment (1 µg per mouse) via subcutaneous administration of recombinant NRG1 containing only the EGF core domain of NRG1-β1 (R&D systems). This form of NRG1 has been shown previously to penetrate the bloodbrain barrier and functionally activate ErbB4 in the cortex (Abe et al., 2011; Sun et al., 2016).

#### Monocular Deprivation

For ocular dominance plasticity experiments in adults (∼P90), one eye lid was closed for 4 days using two mattress sutures (7-0 silk, Ethicon) and checked daily. To produce visual deficits of visual perception, closure of the one eye was maintained for the duration of the normal critical period (∼P18-P32) and was checked daily. If the suture had signs of fraying, a small drop of tissue adhesive (Vetbond) was applied to the suture knot. If an animal was found to have an eye that opened or if the eye was damaged, it was excluded from the experiments. At P32 the eye is re-opened and mice are treated with either saline or subanesthetic ketamine (10mg/kg; s.c.) every other day for three treatments.

#### Intrinsic Signal Optical Imaging

Monocular deprivation was initiated at ∼P90. After 4 days, the sutured eyelid was opened and the skull over the visual cortex was exposed and then covered with agarose (1.5% w/v in 1X PBS) and a coverslip. The agarose and edges of the coverslip were sealed using sterile ophthalmic ointment (Rugby) to prevent drying. A recording session was then initiated.

The primary visual cortex was mapped using intrinsic signal optical imaging through the intact skull as has been described previously (Davis et al., 2015). A custom-designed macroscope (Nikon 135 x 50 mm lenses) equipped with a Dalsa 1M30 CCD camera was used to collect 512 x 512 pixel images sampled at 7.5 Hz (2.2 x 2.2 mm image area). The surface vasculature was first visualized using a 530 nm LED light and the intrinsic signal was then visualized with a 617 nm LED light (Quadica). The camera was focused ∼600 µM beneath the pia surface. Custom written Matlab (Mathworks) code was used to acquire images and to stream them to disk. Visual stimuli (described below) were presented and response data was collected in 5-min session for a total of 3-4 imaging sessions per eye.

Responses to stimulation of the contralateral and ipsilateral eye were recorded from the visual cortex. A visual stimulus was periodically presented and swept within −18 to 26 degrees of the visual field elevation. This stimulus was created by multiplying a band limited (<0.05 cyc/degree; >2 Hz) spatiotemporal noise movie with a one dimensional Gaussian spatial mask (20 degrees) that was phase modulated at 0.1 Hz. Stimuli were presented on an Acer V193 monitor (30 x 37 cm, 60 Hz refresh rate, 20 cd/m^2^ mean luminance). Stimuli were alternatively presented for 5 minutes to each eye. Recordings were made at 0 degrees and 180 degrees. Maps of amplitude and phase of cortical responses were extracted from optical imaging movies via Fourier analysis of each pixel at the frequency of stimulus repetition (0.1 Hz) using custom written Matlab code. Overall map amplitude was computed by taking the maximum of the Fourier amplitude map smoothed with a 5 x 5 Gaussian kernel. Ocular dominance index (ODI) was computed as ODI = (C-I)/(C+I) where C and I are the averaged map amplitudes calculated for contralateral and ipsilateral visual stimulation respectively.

#### Visual Water Maze Task

Visual acuity was assessed using a forced choice, two alternative discrimination task in a visual water maze (Prusky et al., 2000). By testing each eye independently, visual acuity in that eye was determined. A post was surgically implanted onto the skull of the mouse using dental cement. The post allowed for the attachment of a removable custom-made eye occlude to test each eye independently. A week after surgery, mice were trained and then tested. Mice were motivated to swim toward a hidden platform whose location was indicated by the vertical sine wave gratings displayed on a screen at a spatial frequency of 0.063 cpd when viewed from the water maze choice plane. Mice continued training until they reached 90%-100% correct performance levels. Testing was then completed one eye at a time. During testing, the stimulus was increased 0.032 cpd when a mouse achieved at least 70% correct over 10 trials. Mice were required to complete 3 consecutive trials at frequencies under 0.28 cpd in order to automatically advance. For frequencies over 0.28 cpd mice were required to complete 5 consecutive trials in order to automatically advance. Mice were required to complete a block of 10 if any error was made, and the 70% rule was enforced. If performance fell below the 70% correct rate the stimulus was decreased by at least 0.096 cpd. Testing took roughly 200-250 trials per mouse to complete. The threshold of acuity was determined by the spatial frequency corresponding to the 70% value of sigmoidal fit to performance data.

### Translating Ribosome Affinity Purification (TRAP)

Purification of polysomally bound mRNA from visual cortical lysate was performed as described with modifications (Zhou et al., 2013). Briefly, visual cortex was dissected in ice-cold ACSF (in mM: 126 NaCl, 2.5 KCl, 26 NaHCO_3_, 2 CaCl_2_, 2 MgCl_2_, 1.25 NaH_2_PO_4_, and 10 glucose).

Pooled visual cortex from two to five mice was grinded to powder on dry ice, followed with sonication for 5 seconds in ice-cold lysis buffer [20 mM HEPES (pH 7.4), 150 mM KCl, 5 mM MgCl_2_, 0.5 mM dithiothreitol, 100 μg/ml cycloheximide (Sigma-Aldrich), protease inhibitors (Roche) and 40 U/mL recombinant RNase inhibitor (Promega, Madison, WI)]. Homogenates were centrifuged for 10 minutes at 2,000x g, 4 °C, to pellet nuclei and large cell debris, and NP40 (Invitrogen, Carlsbad, CA) and DHPC (Avanti Polar Lipids, Alabaster, Alabama) were added to the supernatant at final concentrations of 1% (vol/vol) and 30 mM, respectively. After incubation on ice for 5 minutes, the lysate was centrifuged for 10 minutes at 20,000x g to pellet insoluble material. Two mouse monoclonal anti-GFP antibodies, Htz-GFP19C8 and Htz-GFP19F7 (50 μg each, Memorial Sloan-Kettering Monoclonal Antibody Facility, New York, NY) were added to bind to 375 μL protein G Dyna magnetic beads (Invitrogen). Alternatively, 300 μL of the streptavidin Myone T1 dynabeads were binding to 120 μg biotinylated protein L first, then followed with two mouse monoclonal anti-GFP antibodies incubation. After being washed twice with the polysome extraction buffer, the beads were then added to the cell-lysate supernatant, and the mixture was incubated at 4 °C with end-over-end rotation for 30 minutes. The beads were subsequently collected on a magnetic rack and washed four times with high-salt polysome wash buffer [20 mM HEPES (pH 7.4), 350 mM KCl, 5 mM MgCl_2_, 1% NP-40, 0.5 mM dithiothreitol and 100 μg/mL cycloheximide]. RNA was eluted from the beads by incubating beads in RLT buffer (RNeasy Micro Kit, Qiagen, Venlo, Netherlands) with β-mercaptoethanol (10 μL/mL) for 5 minutes at room temperature. Eluted RNA was purified using RNeasy Micro Kit (Qiagen) per the manufacturer’s instructions including in-column DNase digestion. Immunoprecipitated RNA yield for each sample was approximately 20 ng/μL.

For PV-Cre; fsTRAP experiments, both cortices (V1) from 5 mice were pooled together to generate n=1 before mRNA extraction. For Emx1-Cre; fsTRAP experiments both cortices (V1) from 2 mice were pooled together to generate n=1 before mRNA extraction.

### Quantitative Real-Time Polymerse Chain Reaction (qPCR)

Purified RNA (30ng) was converted to cDNA using Superscript® III reverse transcriptase (Thermo Fisher Scientific, Waltham, MA, USA) according to the manufacturer’s instructions. Quantitative changes in cDNA levels were determined by real-time PCR using the Power SYBR Green Master Mix (Thermo Fisher Scientific, Waltham, MA, USA), using primers at a concentration of 500nM. Primers were custom designed for mouse NRG1, ErbB4, and for the endogenous control GAPDH. PCR was carried out for 2 min 50 °C, 5 min 95 °C, 40 cycles (15 seconds for 95°C, 30 seconds for 50°C), followed by a melt curve. Technical triplicates were used. GAPDH was used to normalize gene expression and data presented at mean ± SEM. Cycling and quantitation were performed on a ViiA™ 7 Real-Time PCR System instrument (Thermo Fisher Scientific, Waltham, MA, USA) using the ViiA 7 software v1.2. The following primers were used: NRG1-F GCAAGTGCCCAAATGAGTTTAC; NRG1-R GCTCCTCCGCTTCCATAAAT; ErbB4-F CATGGCCTTCCAACATGACTCTGG; ErbB4-R GGCAGTGATTTTCTGTGGGTCCC; GAPDH-F TGCCAACATCACCATTGTTGA; GAPDH-R TGCCAACATCACCATTGTTGA.

### Immunohistochemistry

For immunochemical staining experiments, animals were first deeply anesthetized with Uthasol (sodium pentobarbital, 100 mg/kg, i.p.) and were then perfused transcardially with 5mL of 1X phosphate buffered saline (PBS, pH 7.3–7.4), followed by 20 mL 1X PBS containing 4% paraformaldehyde (PFA) and phosphatase inhibitor (PhosSTOP, 1 tablet for 20 mL, Roche, Switzerland). Brains were removed and maintained in 4% PFA for 24 hours, and then transferred to 30% sucrose in 1X PBS for 24 hours. Then, using a freezing microtome (Leica SM2010R, Germany), coronal sections of the brain were taken at a 30 µm thickness. Mouse V1 coronal sections from bregma −3.40 to −3.80 mm were used for immunohistochemical staining and analysis.

Free floating sections were rinsed 5 times with 1X PBS, and incubated in a blocking solution for 1 hour at room temperature on a shaker. The blocking solution contained 5% normal donkey serum and 0.25% Triton X in 1X PBS. Sections were then incubated with the primary antibody diluted in blocking solution for 36 hours at 4 °C. After incubation with the primary antibody, brain sections were rinsed thoroughly with 1X PBS, and then incubated with the secondary antibody diluted in blocking solution for 2 hours at room temperature. After the secondary antibody was rinsed off, sections were counterstained with 10 µM 4’-6-diamidino-2-phenylindole (DAPI; Sigma-Aldrich, St. Louis, MO) for 5 minutes to help distinguish cortical laminar structure and neuronal nuclei. Lastly, sections were rinsed and then mounted on microscope slides. Sections were cover-slipped with Vectashield mounting medium (H-1000, Vector, Burlingame, CA). To identify PV positive neurons in PV-Cre; fsTRAP mice, the primary goat anti-PV antibody (PVG-213, Swant, Switzerland; RRID:AB_10000345; 1:1000) and a Cy3-conjugated donkey anti-goat antibody (Jackson ImmunoResearch, 1:200) were used. To identify GABA positive neurons in Emx1-Cre; fsTRAP mice, the primary rabbit anti-GABA antibody (SigmaAldrich, 1:1000) and a Cy3-conjugated donkey anti-rabbit antibody (Jackson ImmunoResearch, 1:200) was used. To identify pCREB positive neurons, the primary rabbit anti-pCREB antibody (Cell Signaling, 1:1000) was used.

Immunostained sections were examined, and 10X and 40X image stacks were acquired using a confocal microscope (LSM 780, Carl Zeiss Microscopy,Germany). Image tiles, overlaying, maximum projections, and subset z-stack selections were performed using the Zeiss image processing software. For fluorescent imaging, all sections of a staining series were acquired using the same settings (laser power, pinhole size, line scans), and data images were digitally processed identically.

### Electrophysiology and Laser Scanning Photostimulation

Coronal sections (400uM thick) of V1 were cut from mouse with a vibratome (VT1200S, Leica Biosystems, Buffalo Grove, IL) in sucrose containing ACSF (85 mM NaCl, 75 mM sucrose, 2.5 mM KCl, 25 mM glucose, 1.25 mM NaH_2_PO_4_, 4 mM MgCl_2_, 0.5 mM CaCl_2_, and 24 mM NaHCO_3_). Slices were incubated for at least 30 minutes in sucrose-containing ACSF at 32°C before being transferred into slice-recording chambers with standard ACSF (126 mM NaCl, 2.5 mM KCl, 26 mM NaHCO_3_, 2 mM CaCl_2_, 2 mM MgCl_2_, 1.25 mM NaH_2_PO_4_, and 10 mM glucose). Throughout the cutting, incubation and recording, the solutions were continuously supplied with 95% O_2_–5% CO_2_.

We have previously described our methods for electrophysiological recording, imaging, and photostimulation in detail, including the definitions of all reported parameters (Shi et al., 2010; Xu et al., 2010). For our more recent publications using these same methods see (Kuhlman et al., 2013; Sun et al., 2014; Xu et al., 2016; Xu et al., 2010). Briefly, whole-cell recordings were performed in oxygenated ACSF at room temperature under a differential interference contrast (DIC)/fluorescent Olympus microscope (BX51WI). Oxygenated ACSF was fed into the slice recording chamber through a custom-designed flow system, driven by pressurized 95% O_2_–5% CO_2_ (3 PSI) with a perfusion flow rate of about 2 mL/minute. Slices were first carefully examined under a 4x objective for targeting either L2/3 PV interneurons that express red fluorescent protein (RFP) or tdTomato or pyramidal neurons within the binocular regions of mouse V1, using landmarks defined in the previous study (Antonini et al., 1999). To perform whole-cell recordings, neurons were visualized at high magnification (60x objective, 0.9 NA; LUMPlanFl/IR, Olympus America Inc). The cell bodies of recorded neurons were at least 50uM below the surface of the slice. Patch pipettes (4–6 MΩ resistance) made of borosilicate glass were filled with an internal solution containing 126 mM K-gluconate, 4 mM KCl, 10 mM HEPES, 4 mM ATP-Mg, 0.3 mM GTP-Na, and 10 mM phosphocreatine (pH 7.2, 300 mOsm) when measuring excitatory postsynaptic currents (EPSCs) and action potentials (APs). In separate experiments, a cesium-based internal solution containing 130 mM CsOH, 130 mM D-gluconic acid, 0.2 mM EGTA, 2 mM MgCl_2_, 6 mM CsCl, 10 mM HEPES, 2.5 mM ATP-Na, 0.5 mM GTP-Na, and 10 mM phosphocreatine (pH 7.2, 300 mOsm) was used to voltage clamp pyramidal neurons at the excitatory reversal potential (0–5mV) and measure inhibitory postsynaptic currents (IPSCs). The electrodes also contained 0.1% biocytin for post-hoc cell labeling and further morphological labeling. Once stable whole-cell recordings were achieved with good access resistance (<30 MΩ), basic electrophysiological properties were examined through depolarizing and hyperpolarizing current injections. Electrophysiological data were acquired with a Multiclamp 700B amplifier (Molecular Devices), data acquisition boards (models PCI MIO 16E-4 and 6713, National Instruments), and a custom-modified version of Ephus software 5 (Ephus, available at https://openwiki.janelia.org/). Data were digitized at 10 kHz. Any recordings in which the access resistance changed by >20% during the course of the experiment were excluded from analysis.

During photostimulation experiments, the microscope objective was switched from 60x to 4x for Laser scanning photostimulation (LSPS). The same low-power objective lens was used for delivering ultraviolet flash stimuli. Stock solution of MNI-caged-l-glutamate (Tocris Bioscience) was added to 20 mL of circulating ACSF for a concentration of 0.2 mM caged glutamate. The cortical slice image, acquired through the 4x objective, was visualized using a high-resolution digital CCD camera, and this was in turn was used for guiding and registering photostimulation sites. A laser unit (DPSS Lasers) was used to generate 355 nm UV laser pulses for glutamate uncaging. Short pulses of laser flashes (1 ms, 20 mW) were delivered using an electro-optical modulator and a mechanical shutter. Focal laser spots approximated a Gaussian profile with a diameter of ∼50 μm.

Voltage clamping the recorded neuron allowed determination of sites contributing direct synaptic input. By systematically surveying synaptic inputs from hundreds of different sites across a large region, aggregate synaptic input maps were generated for individual neurons. For our mapping experiments, a standard stimulus grid (16×16 stimulation sites, 65 µm^2^ spacing) was used to tessellate V1 from pia to white matter. The LSPS site spacing was empirically determined to separate adjacent stimulation sites by the smallest predicted distance in which photostimulation differentially activated adjacent neurons. Glutamate uncaging laser pulses were delivered sequentially in a nonraster, nonrandom sequence, following a “shifting-X” pattern designed to avoid revisiting the vicinity of recently stimulated sites (Shepherd et al., 2003). Because glutamate uncaging agnostically activates both excitatory and inhibitory neurons, we empirically determined the excitatory and inhibitory reversal potentials in L2/3 pyramidal cells to properly isolate EPSCs and IPSCs. We voltage clamped PV and pyramidal cells at −70 mV to determine LSPS evoked EPSCs.

Photostimulation induces two kinds of excitatory responses: (1) responses that result from direct activation of the recorded neuron’s glutamate receptors, and (2) synaptically mediated responses (EPSCs) resulting from the suprathreshold activation of presynaptic excitatory neurons. Responses that occur within 10 ms of the laser pulse onset are considered direct; these responses exhibit a distinct waveform and occur immediately after glutamate uncaging. Synaptic currents with such short latencies are not possible because they would have to occur before the generation of action potentials in photostimulated neurons. Therefore, direct responses are excluded from local synaptic input analysis, but they are used to assess glutamate-mediated excitability/responsiveness of recorded neurons. At some locations, synaptic responses over-ride the relatively small direct responses, and these responses are identified and included in synaptic input analysis. For data map analysis, we implement the approach for detection and extraction of photostimulation-evoked postsynaptic current responses described in reference (Shi et al., 2010). LSPS evoked EPSCs are quantified across the 16×16 mapping grid for each cell, and 2 to 4 individual maps are averaged per recorded cell, reducing the likelihood of incorporating noise events in the analysis window. The EPSC input from each stimulation site is the measurement of the sum of individual EPSCs within the analysis window (>10 ms to 160 ms post photostimulation), with the baseline spontaneous response subtracted from the photostimulation response of the same site. The value is normalized with the duration of the analysis window (i.e., 150 ms) and expressed as average integrated amplitudes in picoamperes (pA). The analysis window is used because photostimulated neurons fire most of their action potentials during this time. For the color-coded map displays, data are plotted as the average integrated EPSCs amplitude per pixel location (stimulation site), with the color scale coding input strength. For the group maps obtained across multiple cells, the individual cell maps were first aligned by their slice images using laminar cytoarchitectonic landmarks. Then a new map grid is created to re-sample and average input strength at each site location across cell maps; a smooth version of color-coded map is presented for overall assessments. To further quantitatively compare input strength across cell groups or different conditions, we measure the total ESPC inputs across all map sites (total synaptic input strength) for individual cells.

For the experiments that examine the effects of bath application of NRG1 (R&D systems), MK801 or (2R,6R)-hydroxynorketamine (HNK) (Tocris Bioscience), the reagent(s) are added into the recording solution with specified concentrations. The drug application for 20 minutes was estimated to produce full effects, while the washout of 20-30 minutes was considered to remove the added drug from the recording solution.

### Miniscope Imaging Experiments

The general procedure for surgery has been described previously (Sun et al., 2019). Using the picospritzer setup AAV2-CaMKIIa-GCaMP6f was injected into either bV1. Two weeks later a gradient refractive index (GRIN) lens was implanted at the injection sites for bV1. For bV1 implantation a 1.8mm-diameter circular craniotomy was centered at the injection coordinates. For bV1 implantations, the dura mater was removed above the cortex, but no cortex was removed. The GRIN lens (.25 pitch, 0.55 NA, 1.8mm diameter and 4.31mm in length; Edmund Optics) was slowly lowered with a stereotaxic arm onto the surface of the cortex for bV1. In order to anchor the GRIN lens to the skull, a skull screw was used. The GRIN lens and skull screw were fixed to the skull with cyanoacrylate and dental cement. Kwik-Sil (World Precision Instruments) was used to cover the lens. After 2 weeks a small aluminum baseplate was cemented onto the mouse’s head over the existing dental cement in order to provide the miniscope a place for attachment to the animal. The miniscope was then fitted to the baseplate and locked in a position so that GCaMP6f expressing neurons were in focus.

For behavioral experiments involving bV1, mice were first handled for 10 minutes per day for a week in order to allow the animals to become comfortable with the behavioral experimenter. Then animals were habituated to the behavioral arena for 10 minutes per day for 2 days without a miniscope. Behavior was performed in a quiet designated behavioral room under low lighting conditions (<20 lux). The behavioral arena was 35cm x 25 cm in size with bedding which was changed between animals. Animals were then habituated to the behavioral arena with the miniscope fixed onto their head while they behaved freely for 10 minutes per day for 3 days, so that animals were acclimated and habituated to the behavioral experimenter, arena, and miniscope gear. After preparations were completed, *in vivo* GCaMP6-based calcium imaging of population bV1 neurons was performed in awake freely behaving mice. Behavioral recording were made by tracking the location of the LED on the miniscope through the experimental sessions. Sessions were 5-10 minutes in length.

Please refer to www.miniscope.org/ for technical details about miniscope construction. Miniscopes are head-mounted and have a mass of ∼3 grams. The miniscope has a 700 µm x 450 µm field of view with a resolution of 752 pixels x 480 pixels (∼1 µM per pixel). The data is transmitted over a Super Speed USB to a PC running custom DAQ software after the DAQ electronics package the data to comply with the USB video class (UVC) protocol. The DAQ software uses Open Computer Vision (OpenCV) libraries for image acquisition and is written in C++. Images are acquired at ∼30 frames per second (fps) and then recorded to uncompressed .avi files. The DAQ software also recorded the animal’s behavior with a high definition webcam (Logitech) at ∼30 fps, and time stamped both video streams for alignment offline.

The individual sessions of miniscope videos were concatenated and then down-sampled by a factor of 2 using NIH ImageJ software, and then videos were motion corrected using the NoRMCorre MATLAB package. Subsequent analysis was performed using custom MATLAB scripts by adopting the method of extended constrained non-negative matrix factorization for endoscopic data (CNMF-E) (Zhou et al., 2018). The CNMF-E is based on the CNMF framework and enables simultaneously denoising, deconvolving and demixing calcium imaging data. This enables the extraction of individual neuron calcium activity from concatenated videos. The key features of CNMF-E include the separation of single neuron signal from a large rapid fluctuating background signal that has low spatial-frequency structure. CNMF-E extracts the spatial footprints of neurons and their associated temporal calcium activity after iteratively solving the constrained matrix factorization problem. See more detailed information in our published work (Sun et al., 2019). To assess single cell Ca++ signal responses, we measured peak calcium event amplitudes and summed calcium event amplitudes from extracted neurons (Figure S7).

### Visually Evoked Potential (VEP) Recording

The general procedure for surgery has been described previously (Kaplan et al., 2016). A steel head bar was affixed to the skull anterior to bregma using cyanoacrylate glue. Burr holes (< 0.5 mm) were then drilled into the skull over binocular V1 (∼3.1 mm lateral of lambda) over the region identified as binocular visual cortex using intrinsic signal optical imaging. Tapered tungsten recording electrodes (FHC, Bowdoinham, ME, US), 50 μm in diameter at their widest point, were implanted in each hemisphere, ∼350 μm below the visual cortical surface into deep L2/3. Silver wire (A-M systems, Sequim, WA, US) reference electrodes were implanted over prefrontal cortex. Mice were allowed to recover for at least 2 weeks prior to experimentation.

Visual stimuli were generated using software (http://bearlab-s1.mit.edu/supp6/PlxStimOne_Opto.zip) written and shared by Jeffrey Gavornik in either C++ for interaction with a VSG2/2 card (Cambridge Research systems, Kent, U.K.) or Matlab (MathWorks, Natick, MA, U.S.) using the PsychToolbox extension (http://psychtoolbox.org) to control stimulus drawing and timing. The display was positioned 20 cm in front of the mouse and centered, thereby occupying 92° × 66° of the visual field. Visual stimuli consisted of full-field bright to dark screen reversal at a frequency of 2 Hz. Bright screen stimulus luminance was X cd/m^2^. Experiments were fully automated and each session consisted of 400 phase reversals and included two 30-s intervals during which the screen was gray for spontaneous response data collection.

VEP recordings were conducted in awake, head-restrained mice. Prior to recording, mice were habituated to the restraint apparatus by sitting in situ in front of a gray screen for a 15-minute session on each of three consecutive days. On stimulus presentation days, VEP magnitudes were quantified by measuring trough-baseline response magnitude averaged over all phase reversals. Recordings were amplified and digitized using the Neuralynx electrode adaptor interface board (EIB-16) (Neuralynx., Bozeman, MT), and the Intan headstage preamplifier (RHD-2132) and Intan RHD2000 USB Interface Board (Intan, Los Angeles, CA). A recording channel was dedicated to recording VEPs from bV1 in each implanted hemisphere and another recording channel was reserved for collecting reference information. Local field potentials were recorded from V1 with 20 kHz sampling rate and a 250 Hz low-pass filter, and a notch filter (59-61 Hz). Data was extracted from the binary storage files generated by the Intan RHD2000 Evaluation software package and analyzed using custom software (http://bearlab- s1.mit.edu/supp6/VEP_Analysis.zip) written in C++, Matlab and Labview. VEPs were averaged across all stimulation reversals within a session of 400 trials, and trough-baseline differences were measured within a 100-millisecond period from bright screen start.

### Two-Photon Calcium Imaging

The general procedure for surgery has been described previously (Davis et al., 2015; Kuhlman et al., 2013). After the skull was exposed, it was dried and covered by a thin layer of Krazy glue. After the Krazy glue dried (∼15 min), it provided a stable and solid surface on which to affix an aluminum headplate with dental acrylic. The headplate was affixed to the skull and the margins sealed with dental acrylic to prevent infections. A 4-mm-diameter region of skull overlying the visual cortex was removed. The craniotomy opened at center coordinates 3mm lateral, and 1.7mm anterior to the lambda suture. Care was taken to leave the dura intact. A sterile, 4-mm-diameter cover glass was then placed directly on the dura and sealed at its edges with Krazy glue. When dry, the edges of the cover glass were further sealed with dental acrylic. At the end of the surgery, all exposed skull and wound margins were sealed with VetBond and dental acrylic. Mice were then removed from the stereotaxic apparatus, given a subcutaneous bolus of warm sterile saline and carprofen, and allowed the mouse to recover on the heating pad. When fully alert, they were placed back in their home cages, and gave carprofen daily for at least 3 days. Typically, the mouse was permitted to recover for at least seven days and conditioned to the head restraint and running wheel for several days, 20-minutes per day before mapping. And wait for 10-14 days after surgery to perform imaging.

Two-photon calcium imaging was performed using a resonant scanning, two-photon microscope (Neurolabware, Los Angeles, CA) controlled by Scanbox acquisition software (Scanbox, Los Angeles, CA). The light source was a Coherent Chameleon Ultra II laser (Coherent Inc, Santa Clara, CA) running at 920nm. The objective was an 16X water immersion lens (Nikon, 0.8NA, 3mm working distance). Usually we used the bidirectional scanning mode for the imaging. the microscope frame rate was 16.05 Hz (494 lines with a resonant mirror at 8kHz). The imaging field covered ∼700 µm × 500 μm. Eye movements and pupil size were recorded via a Dalsa Genie M1280 camera (Teledyne Dalsa, Ontario, Canada) fitted with a UV cut-off filter (wavelength range 350 – 2200 nm). Images were captured at layer 2/3 (150-300um below pia). During imaging a substantial amount of light exits from the brain through the pupil. Thus, no additional illumination was required to image the pupil. Mice were head-fixed on a spherical treadmill on which they had been trained to balance and run. Movement of the spherical treadmill was recorded via a ball bearing option tracker (H5, US digital). Both locomotion and eye movement data were synchronized to the microscope frames. We used the wavelength 920nm to image GCaMP6s activities.

Custom-written Matlab pipelines based on CaImAn (Zhou et al., 2018), which is an open-source library for calcium imaging data analysis, were used to remove motion artifact, extract neurons and neural activity indentation and perform analyses. The change in fluorescence intensity relative to the resting fluorescence intensity (ΔF/F) was used to represent the calcium activities. The average calcium event rate, and average event amplitude were measured for 15 min spontaneous response recording while animals viewed gray screen.

### GCaMP6-based Epi-fluorescence Retinotopic Mapping

Prior to two-photon calcium imaging of binocular visual cortical responses, retinotopic maps were acquired by epi-fluorescence imaging in 4X/0.16 objective. The surface vasculature and GCaMP6s signal were visualized from White Mounted LEDs (400 - 700 nm) (MCWHLP1, Thorlabs), via a bandpass filter (482/18nm, Thorlabs) and fluorescence was detected by a sCMOS camera (PCO.edge 4.2 LT) via a 520/28nm bandpass filter (Thorlabs). The focal plane of the microscope was at 300-400 µm below the surface vasculature. During retinotopic mapping, the head was restrained via the implanted headplate and eyes were on a horizontal plane. Visual stimuli were generated in real-time with Processing using OpenGL shaders. The stimulation was displayed on a 27’’ LCD Monitor (Asus vg 278) refreshed at 60 Hz, placed 20 cm perpendicular from the mouse midline. The monitor subtending approximately −56 to 56 degrees in azimuth and −40 to 40 degrees in altitude. A TTL pulse was generated with an Arduino at each stimulus transition and the pulse was recorded by the microscope and time-stamped which used for synchronizing between visual stimulation and imaging data.

Retinotopic maps were generated by sweeping a bar across the monitor (Kalatsky and Stryker, 2003). The bar contained black and white checkerboard pattern on it, with checker size 4 degree. It was swept across the screen 8 times in each of four cardinal directions with a period of 10s. There is a 10s interval of blank gray screen was inserted between sweeps. Azimuth and elevation retinotopic maps were used to generate a visual field sign maps (Garrett et al., 2014). Binocular V1 was confined to regions adjacent to the intersection of the horizontal and vertical meridians at the border of V1 and LM. *In vivo* two-photon imaging was in the binocular area of V1 identified by retinotopic mapping.

### Statistical Analysis

All data are reported as mean ± SEM. For statistical comparisons between groups, data were checked for normality distribution and equal variance. When comparing two independent groups, normally distributed data were analyzed using a Student’s t test. In the case data were not normally distributed, a Mann-Whitney U test was used. In the case more than two groups were compared and data were normally distributed, a one-way ANOVA was performed and followed by post hoc comparisons when justified. In other cases, a non-parametric Kruskal Wallis test was used and followed by comparisons with Mann-Whitney U tests. A p-value (p < 0.05) was considered statistically significant.

## Acknowledgements

This work was supported by US National Institutes of Health (NIH) grants R01EY028212, R01EY027407, R01 MH105427, R35GM127102 and R01EY029490.

---

**Supplementary Figure 1.**
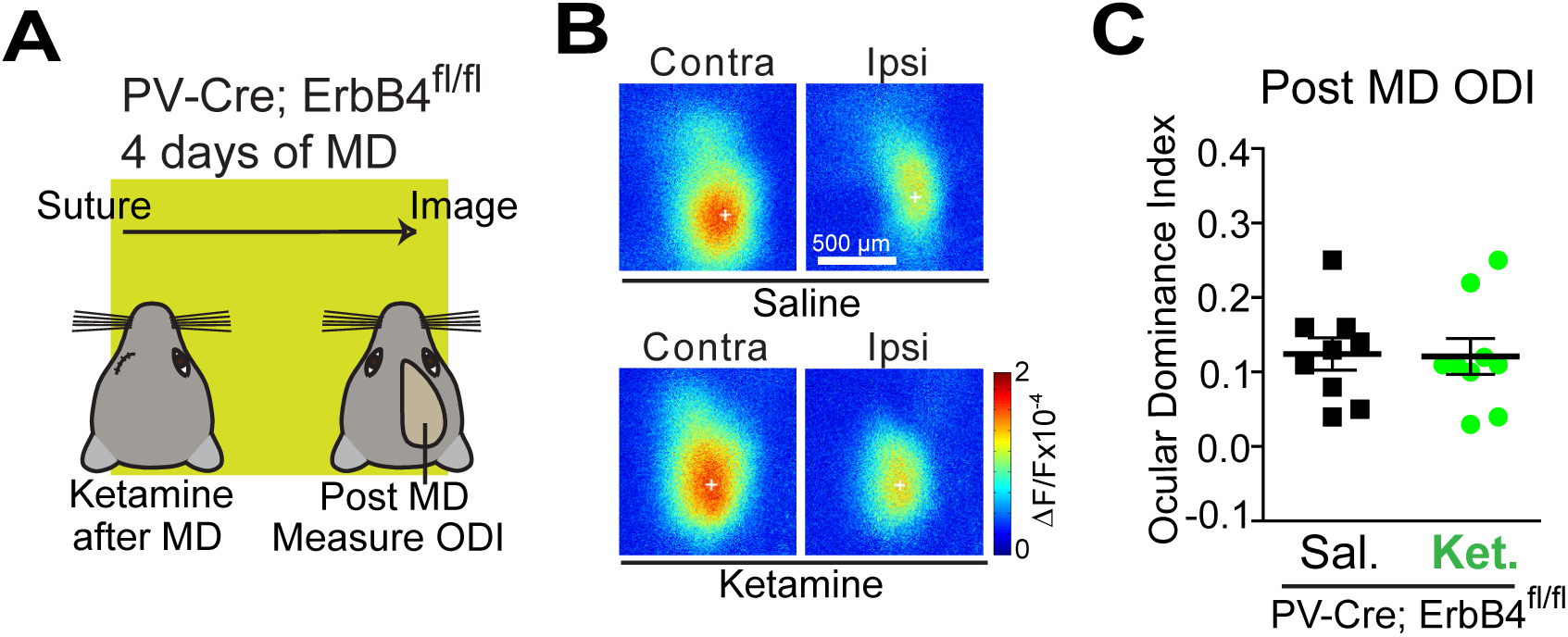
Ketamine does not induce adult ocular dominance plasticity in PV ErbB4 null mice. (A) Illustration of the experimental paradigm for assessing ocular dominance plasticity in adult (P90-120) mice, and the timeline of the experimental protocol. Immediately after monocular deprivation (MD), saline or ketamine treatments are given, and 4 days later the ocular dominance index (ODI) is measured after eyelid suture removal by measuring intrinsic signal responses of visual cortex to contralateral versus ipsilateral eye stimulation. (B) Representative contralateral and ipsilateral cortical response maps are shown from a saline treated PV ErbB4 null animal (top) and a ketamine treated PV ErbB4 null animal (bottom) at 4 days after treatment. (C) ODI does not significantly change in ketamine treated adult PV ErbB4 null animals (n=9, green) as compared to control saline treated animals (n=9, black).

**Supplementary Figure 2.**
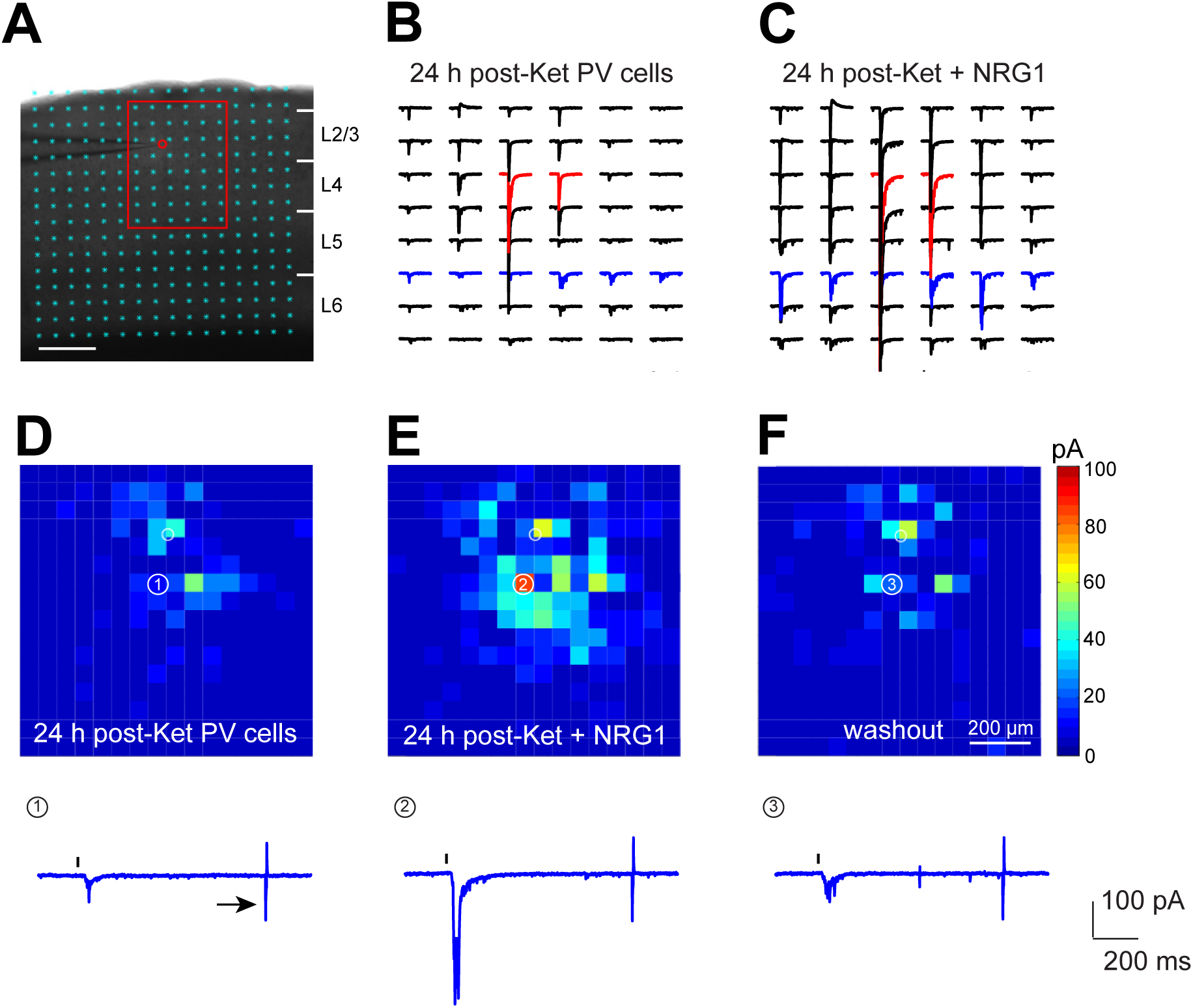
Laser scanning photostimulation (LSPS) allows for quantitative and extensive mapping of local excitatory synaptic inputs to PV neurons. LSPS allows for extensive and quantitative analysis of synaptic inputs to recorded PV cells from laminar circuits in a relatively large cortical region. (A) PV/fast-spiking (FS) cells are targeted by tdTomato expression in PV-Cre; Ai9 mouse V1 slices. A V1 slice image superimposed with photostimulation sites (cyan circles) spaced at 60 μm x 60 μm. The red circle indicates the tip of a recording electrode and the cell body location of a recorded L2/3 PV interneuron from a mouse treated with ketamine. Scale bar = 200 μm. (B-C) The plot of excitatory postsynaptic current (EPSC) responses from the recorded PV cell at the selected sites within the region shown by the red rectangle in (A) before and after bath application of exogenous recombinant NRG1 in a V1 slice from a mouse treated with ketamine. (D-F) Representative example of bath NRG1 enhancement of excitatory synaptic inputs to a ketamine treated PV cell. Data maps were obtained before (D), during (E) and after washout (F) of bath applied NRG1. The spatial scale in (F) indicates 200 μm. The color scale (F) indicates average integrated input strength at individual map sites. The warmer color indicates stronger input strength. The white numbered circle indicates the cell body location of the ketamine treated PV neuron. Each map site (color pixel) is spaced at 60 μm x 60 μm. (Below D-F) Synaptic input responses at the specified, numbered sites. The response traces are plotted for 1200 ms, with 200 ms baselines before a 1ms photostimulation (black ticks above the traces). Current injection responses (5 pA, 5 ms; pointed by the arrow) allow for the monitoring of access resistance during the mapping experiment. Any experiment in which the access resistance changes by >20% during the course of the experiment were excluded from the analysis.

**Supplementary Figure 3.**
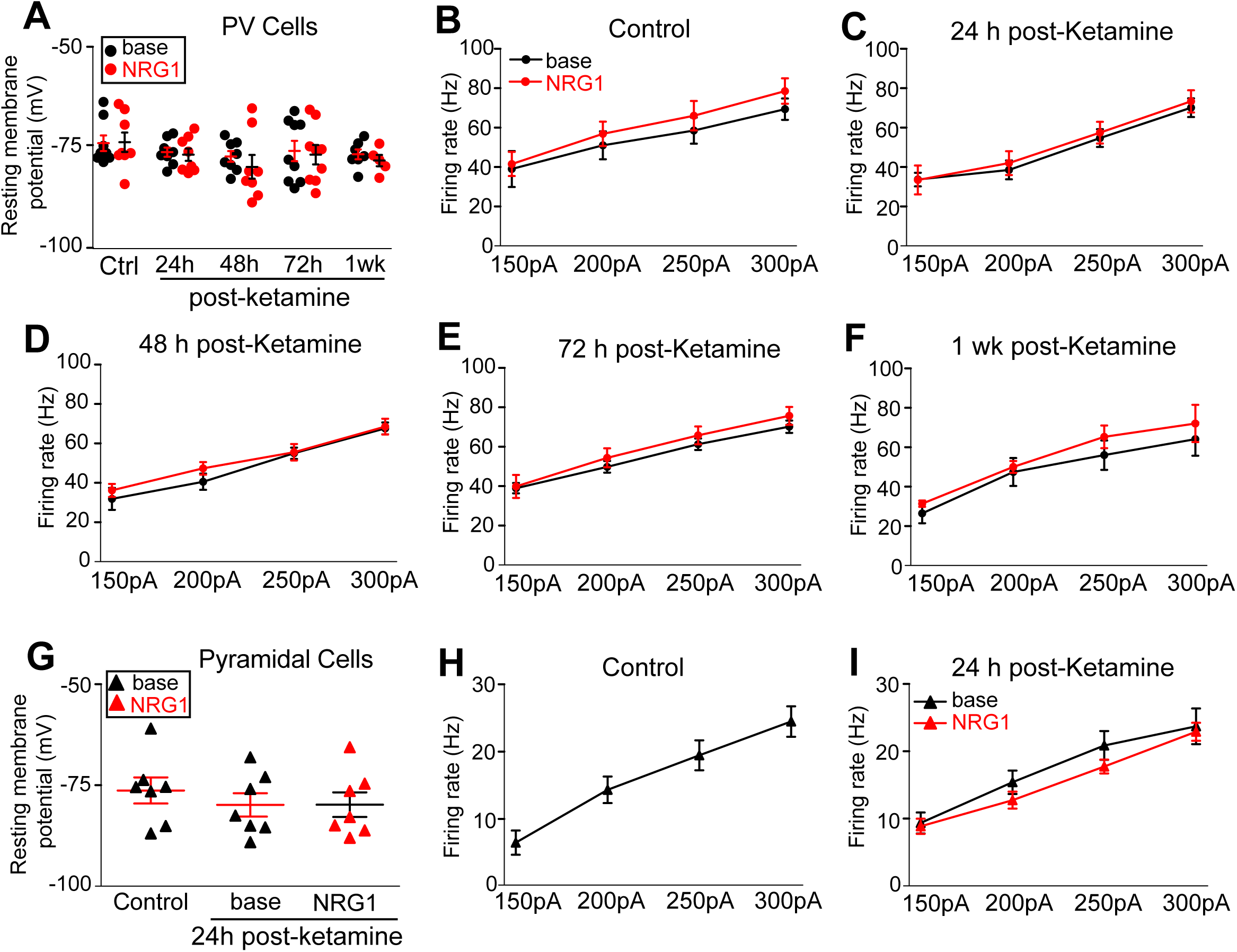
Resting membrane potentials, intrinsic membrane excitability and firing rates of PV neurons or excitatory neurons do not change after ketamine treatment and with NRG1 bath application. (A) Resting membrane potentials of PV cells of control, and 24, 48, 72 hours and 1 week after ketamine treatment do not differ, and they are not changed by bath application of exogenous recombinant NRG1. (B-F) Plots of the overall relationship between PV cell firing rates and current injection strengths at 24 hours (C), 48 hours (D), 72 hours (E) or 1 week (F) post-ketamine treatment as compared to controls (B). The data values are represented as means ± SEM and at each condition [controls (n = 8 cells), 24h (n = 8), 48h (n = 8), 72h (n = 9) and 1wk (n = 7)]. Resting membrane potentials and firing rates were determined before and after bath application of NRG1 (A-F). (G) Resting membrane potentials of L2/3 pyramidal (PYR) cells of control and 24 hours post-ketamine treatment do not differ, and they are not changed by bath application of NRG1. (H,I) Plots of the overall relationship between PYR cell firing rates and current injection strengths for PYR cells at 24 hours after ketamine treatment as compared to controls (H) [controls (n = 7) and 24h post ketamine (n = 7)].

**Supplementary Figure 4.**
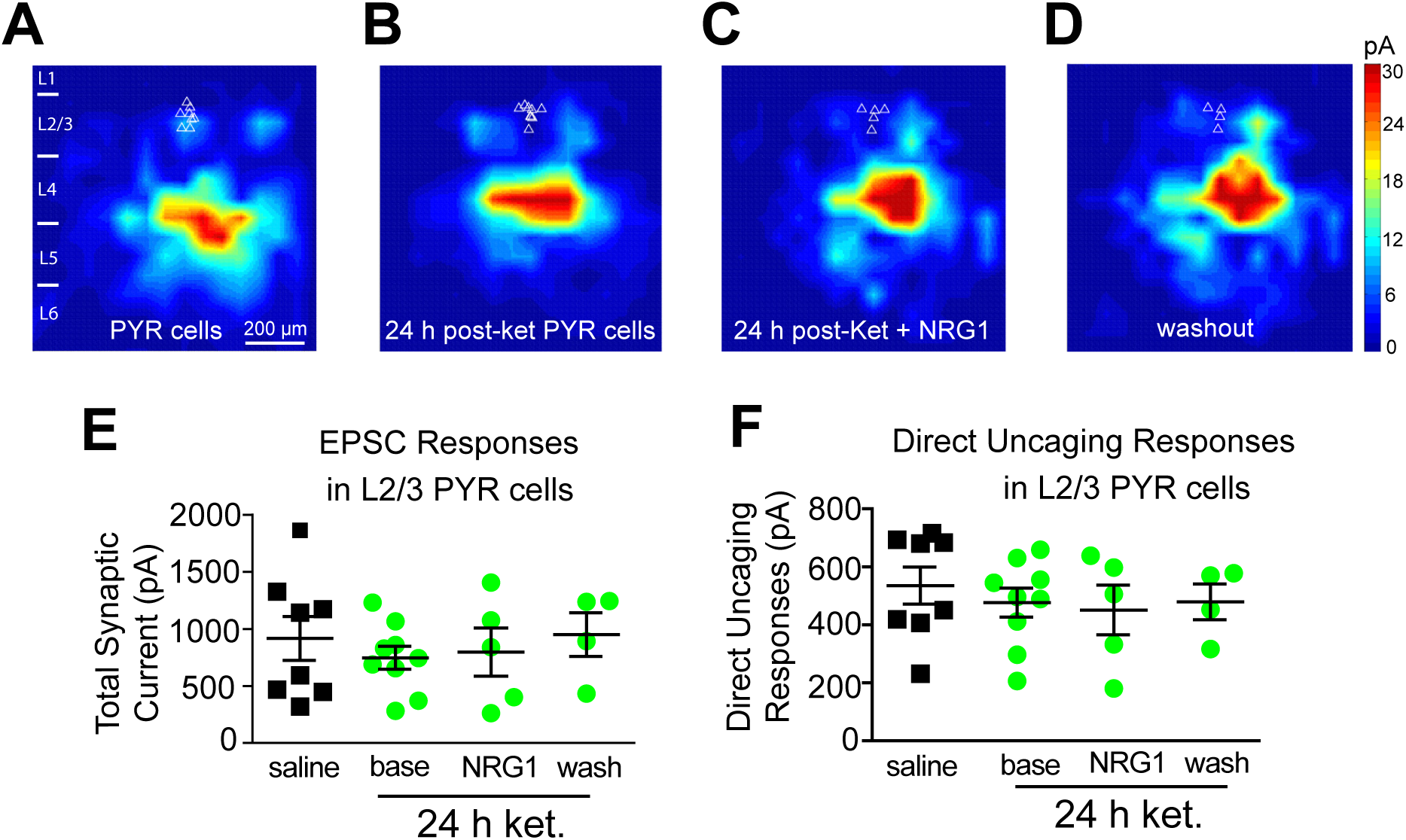
Excitatory inputs to L2/3 pyramidal neurons are not modulated by *in vivo* ketamine treatment. (A-D) LSPS mapping of local excitatory inputs to individually recorded L2/3 pyramidal neurons in the visual cortex of control mice and the mice at 24 hours after ketamine treatment (10 mg/kg; s.c.). Neither ketamine treatment nor an exogenous NRG1 bath application affected the excitatory inputs to L2/3 pyramidal neurons. Representative L2/3 EPSC input maps for the specified conditions are shown. (E-F) Quantitative summary data from A-D show that total synaptic currents and direct uncaging responses of L2/3 pyramidal cells do not differ across conditions.

**Supplementary Figure 5.**
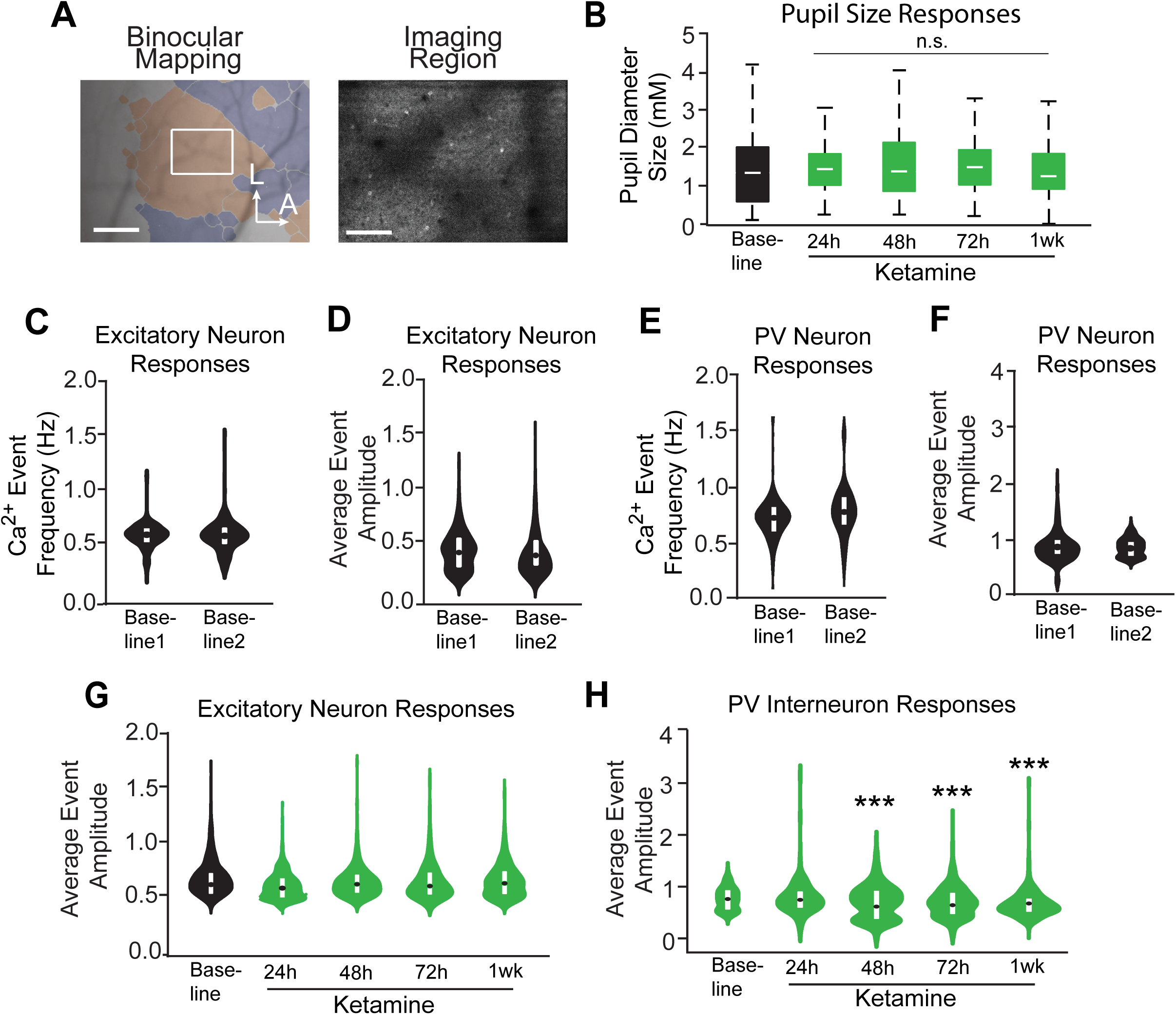
The baseline stability of two-photon calcium recording of excitatory or PV neuron responses. (A, left) Contralateral and ipsilateral eye stimulation is used to determine the binocular visual cortex region via GCaMP6 signal-based binocular mapping with a 4x objective (Olympus) in a 2P rig (Neurolabware). Binocular visual cortex maps are overlaid on images of the existing cortical topography generated with bright field imaging (scale bar = 500 μm). (A, right) The cortical region (bV1) can then be located with a 16x objective (Olympus) for recording of calcium signals during an imaging session. A representative maximum intensity Z-projection is shown (scale bar = 125 μm). (B) Pupil sizes are monitored during imaging sessions (24, 48, 72 hours and 1 week) after ketamine treatment. (C-D) Neither the calcium event frequencies (C) nor the average integrated amplitudes (D) of excitatory neuron responses significantly change between two consecutive days of baseline imaging. (E-F) Neither the calcium event frequencies (E) nor the average event amplitudes (F) of PV neuron responses significantly change between two consecutive days of baseline imaging. (G-H) The average event amplitudes of calcium responses for excitatory (G) or PV (H) neurons 24, 48, 72 hours and 1 week after ketamine treatment. For G-H, *** p < 0.001 (repeated measures ANOVA with Bonferroni’s multiple comparison). Data represent mean ± SEM.

**Supplementary Figure 6.**
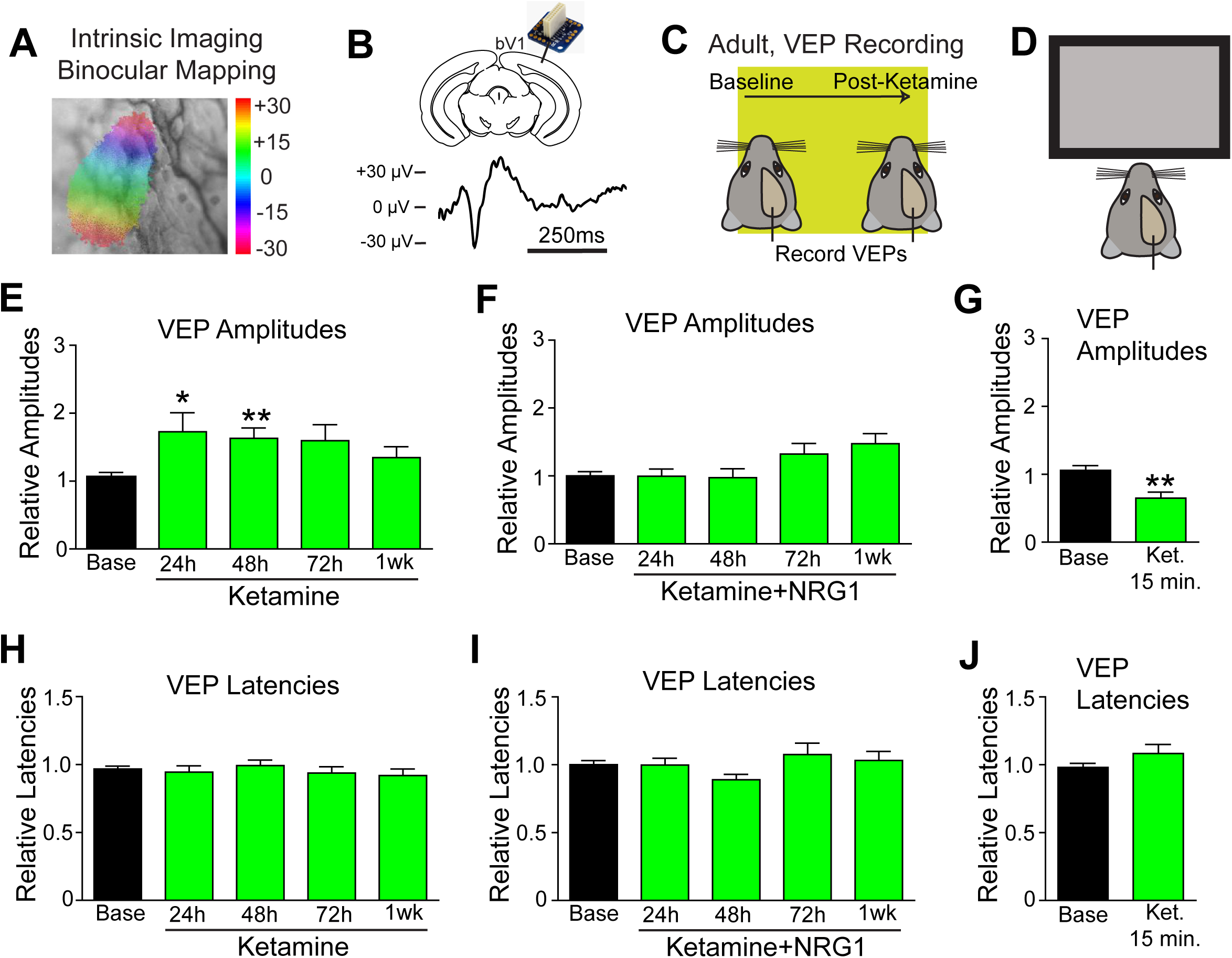
Ketamine increases amplitudes of visual evoked potentials (VEP) in the visual cortex. (A) Intrinsic signal optical imaging responses to contralateral and ipsilateral eye stimulation are used to determine the bV1. (B) Local field potential (LFP) electrodes are then surgically implanted into deep L2/3 in bV1, and are connected to a small interface board mounted on the animal’s skull. (B, bottom) An example VEP trace during baseline recording. (C) Illustration of the timeline of the experiment. Briefly, baseline VEPs are recorded and then ketamine (10mg/kg; s.c.) is administered to adult mice (P90-P120). Subsequent recordings are made at 15 min, 24, 48, 72 h, and 1 week after treatment. (D) Mice are head fixed in front of a high performance monitor in which 400 bright screen visual epochs (500ms) are presented to the mouse per recording session. (E) Ketamine treatment significantly increases the VEP amplitudes 24 and 48 h after treatment (n = 20). (F) *In vivo* exogenous NRG1 treatment (1 μg per mouse, s.c.) at 2 hours before ketamine injection blocks ketamine-induced increases in VEP amplitudes. (G) Within 15 minutes of ketamine treatment, VEP amplitudes are reduced. (H-J) Neither ketamine nor NRG1 treatment has an effect on VEP latencies at the timepoints after treatment.

**Supplementary Figure 7.**
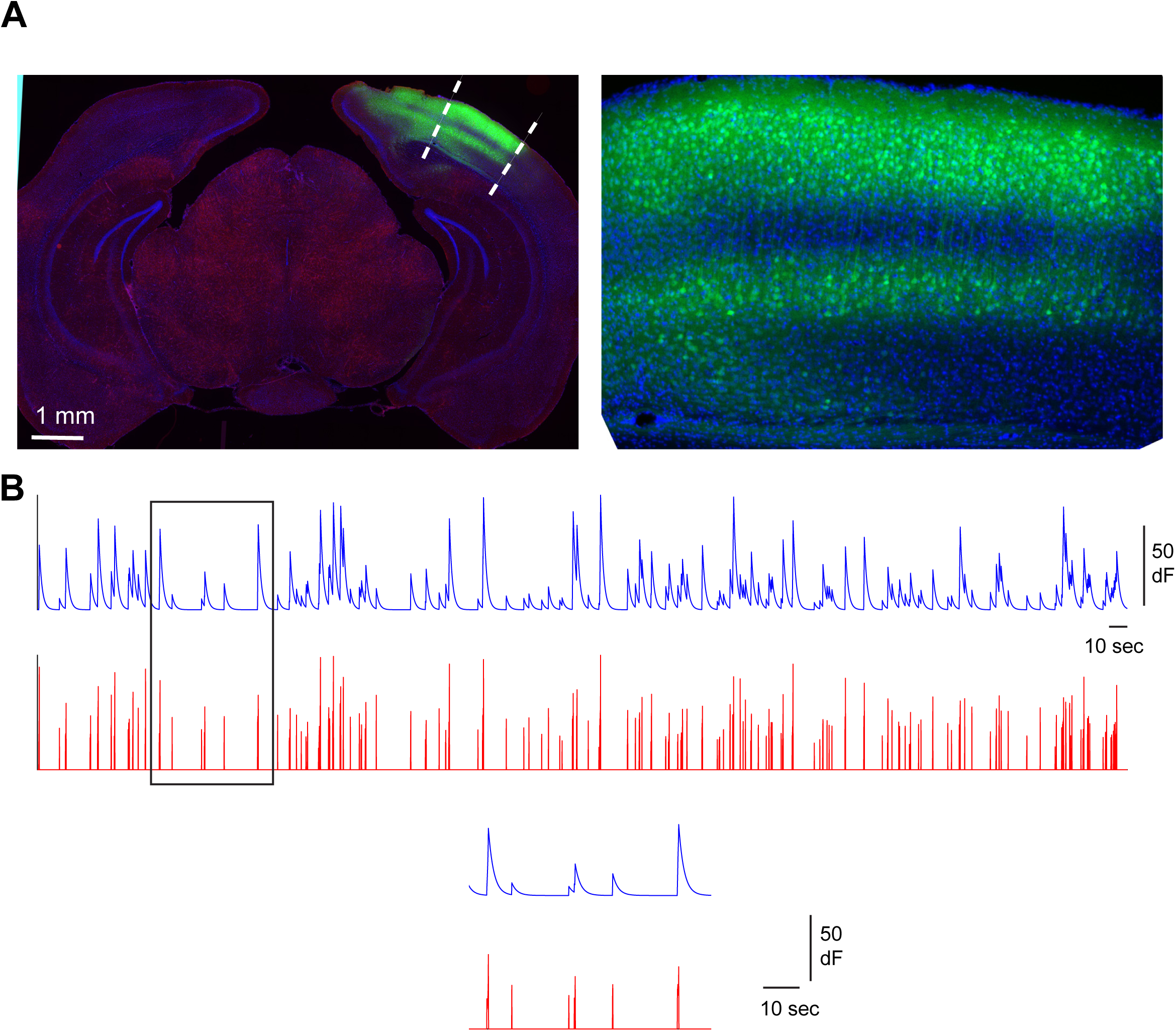
Miniscope GRIN lens placement and illustration of calcium activity extraction. (A) The left image panel shows a coronal brain section from an experiment mouse with the dashed lines indicating the boundaries of a GRIN lens implant. The DAPI staining is blue, and GCaMP6f expression is green. The right panel show an enlarged region of V1 beneath the lens. Overall normal-looking structures and GCaMP6f expression in visual cortex provide support for the imaging of healthy V1 excitatory cells. (B) The upper traces show the temporal trace of calcium signal activity from an example cell from miniscope-based GCaMP6 imaging and the corresponding deconvolved trace (red) is used for inferring spiking activity. Lower traces show the expanded segment of both traces indicated in the black box above. See Methods for more details. We use deconvolved traces for downstream analysis and apply a threshold of 10% of the maxima of peak responses in these traces across all of the sessions to identify effective calcium events. Two-photon calcium imaging data are similarly processed.

**Supplementary Table 1.** Experiments, mouse strains and relevant procedures

■ Intrinsic Signal Imaging (Figure 1; Supplemental Figure 1)
| Mice Used | Treatments | Experiments |
| --- | --- | --- |
| C57BL/6 (N=34) | Saline (s.c) or ketamine (10-50mg/kg; s.c.) | Monocular deprivation and intrinsic optical imaging to determine ODI |
| PV-Cre; ErbB4<sup>fl/fl</sup> (N=19) | Saline (s.c) or ketamine (10-50mg/kg; s.c.) | Monocular deprivation and intrinsic optical imaging to determine ODI |

■ Visual Water Maze Task (Figure 1)
| Mice Used | Treatments | Experiments |
| --- | --- | --- |
| C57BL/6 (N=24) | Saline (s.c) or ketamine (10mg/kg; s.c.) | Monocular deprivation to induce amblyopia visual water maze task during adulthood |

■ IPSC Recording from excitatory pyramidal cells (PYRs) (Figure 2)
| Mice Used | Treatments | Experiments |
| --- | --- | --- |
| C57BL/6 (N=5) | Bath ketamine treatment | PYR IPSC recording after acute bath application of ketamine to cortical slices |
| C57BL/6 (N=5) | Bath MK-801 treatment | PYR IPSC recording after acute bath application of MK-801 to cortical slices |
| C57BL/6 (N=4) | Saline (s.c) or ketamine (10mg/kg; s.c.) (1 hr) | PYR IPSC recording 1 hr after ketamine injection |
| C57BL/6 (N=28) | Saline (s.c) or ketamine (10mg/kg; s.c.)(24, 48, 72 h, 1 wk) | PYR IPSC recording at different times after ketamine injection |
| PV-Cre; ErbB4<sup>fl/fl</sup> (N=10) | Saline (s.c) or ketamine (10mg/kg; s.c.)(24h) | PYR IPSC recording 24 h after ketamine injection |

■ NRG1 and ErbB4 mRNA Quantification (Figure 3)
| Mice Used | Treatments | Experiments |
| --- | --- | --- |
| PV-Cre; fsTRAP (N=3) | none | Perfusion and characterization of EGFP expression in PV cells |
| Emx1-Cre; fsTRAP (N=3) | none | Perfusion and characterization of EGFP expression in excitatory cells |
| PV-Cre; fsTRAP (N=125) | Saline (s.c) or ketamine (10mg/kg; s.c.) (24, 48, 72 h, 1 wk) | Quantification of PV-specific NRG1 and ErbB4 expression by qPCR |
| Emx1-Cre; fsTRAP (N=50) | Saline (s.c) or ketamine (10mg/kg; s.c.)(24, 48, 72 h, 1 wk) | Quantification of excitatory-specific NRG1 and ErbB4 expression by qPCR |

▪pCREB Quantification (Figure 3)
| Mice Used | Treatments | Experiments |
| --- | --- | --- |
| PV-Cre; Ai9 (N=30) | Saline (s.c) or ketamine (10mg/kg; s.c.)(24, 48, 72 h, 1 wk) | Perfusion and measurement of pCREB immunostaining |

▪Intrinsic Properties of PVs and PYRs (Supplementary Figure 3)
| Mice Used | Treatments | Experiments |
| --- | --- | --- |
| PV-Cre; Ai9 (N=20) | Saline (s.c) or ketamine (10mg/kg; s.c.)(24, 48, 72 h, 1 wk) | Resting membrane potential measurements of PV cells after ketamine treatment |
| C57BL/6 (N=8) | Saline (s.c) or ketamine (10mg/kg; s.c.)(24, 48, 72 h, 1 wk) | Resting membrane potential measurements of PYRs after ketamine treatment |

▪EPSC Recordings from PVs and PYRs using LSPS (Figure 4; Supplementary Figure 4)
| Mice Used | Treatments | Experiments |
| --- | --- | --- |
| PV-Cre; Ai9 (N=25) | Saline (s.c) or ketamine (10mg/kg; s.c.)(24, 48, 72 h, 1 wk) | PV EPSC recording at different times after ketamine injection |
| C57BL/6 (N=10) | Saline (s.c) or ketamine (10mg/kg; s.c.) (24 h) | PYR EPSC recording 24 h after ketamine injection |

▪Effect of HNK on PV and PYR cells (Figure 5)
| Mice Used | Treatments | Experiments |
| --- | --- | --- |
| C57BL/6 (N=5) | Saline (s.c) or HNK bath application | PYR IPSC recording at different times after HNK application |
| C57BL/6 (N=5) | Saline (s.c) or HNK (10mg/kg; s.c.)(24h) | PYR IPSC recording 24h after HNK injection |
| PV-Cre; Ai9 (N=5) | Saline (s.c) or HNK (10mg/kg; s.c.)(24h) | PV EPSC recording 24h after HNK injection |

▪2P imaging of PV and Excitatory cells (Figure 6; Supplementary Figure 5)
| Mice Used | Treatments | Experiments |
| --- | --- | --- |
| GCaMP6s transgenic (N=4) | Saline (s.c) or ketamine (10mg/kg; s.c.)(24, 48, 72 h, 1 wk) | 2P recording of excitatory cells at different times after ketamine injection |
| PV-Cre; Ai163 (N=4) | Saline (s.c) or ketamine (10mg/kg; s.c.)(24, 48, 72 h, 1 wk) | 2P recording of PV cells at different times after ketamine injection |

▪VEP recording (Supplementary Figure 6)
| Mice Used | Treatments | Experiments |
| --- | --- | --- |
| C57BL/6 (N=40) | Saline (s.c), ketamine (10mg/kg; s.c.)(24, 48, 72 h, 1 wk), and/or NRG1 (1µg; s.c.) | VEP recording of bV1 at different times after ketamine injection |

▪Miniscope recording (Figure 7; Supplementary Figure 7)
| Mice Used | Treatments | Experiments |
| --- | --- | --- |
| C57BL/6 (N=12) | none | miniscope recording of bV1 in freely behaving animals |
| PV-Cre; ErbB4<sup>fl/fl</sup> (N=13) | none | miniscope recording of bV1 in freely behaving animals |

